# Ecology and engineering to modify the bile acid output of a defined microbial community

**DOI:** 10.64898/2026.05.23.727444

**Authors:** Xianfeng Zeng, Xiandong Meng, Allison M. Weakley, Kelsey E. Jarrett, Steven Higginbottom, Eugenell Mae Lopez, Ashley V. Cabrera, Ira J. Gray, Brian C. DeFelice, Masahiko Terasaki, Rochelle Lai, Madelaine Brearley-Sholto, Aishan Zhao, Kerrigan R. Hall, Mikhail Levia, Jeanette Arreola, Thomas Q. de Aguiar Vallim, Michael A. Fischbach

## Abstract

The bile acid pool, which is synthesized collaboratively by the host and its microbiome, impacts metabolism, immunity, and disease risk. Targeted microbiome interventions could in principle reshape the bile acid pool for therapeutic benefit, but practical strategies remain elusive. In the course of screening a complex defined community for metabolic phenotypes by dropping out individual strains, we observed that several of the single-strain dropout communities had markedly increased deoxycholic and lithocholic acid levels and a larger bile acid pool. In each of these communities, a second strain—*Lactobacillus plantarum—*had bloomed. The bile salt hydrolase activity of *L. plantarum* was necessary and sufficient to expand the size of the bile acid pool. An engineered community in which the *bsh* gene is overexpressed in multiple *Lactobacillus* strains confers on mice increased levels of secondary bile acid levels and a larger pool size. By overexpressing a different pair of bile acid metabolic genes in multiple strains of *Lactobacillus*—7α- and 7β-hydroxysteroid dehydrogenase—we changed the composition of the bile acid pool, enlarging it and redirecting it toward ursodeoxycholic acid. Together, these results demonstrate that fine details of the microbiome’s strain composition can have a substantial effect on bile acid metabolism, and that rational manipulation of the microbiome can alter the size and composition of the bile acid pool.

## INTRODUCTION

The bile acid pool is one of the most concentrated and biologically active sets of metabolites in humans and other animals^1^. Bile acids are synthesized in the liver from cholesterol, stored in the gallbladder, and secreted into the small intestine upon feeding^2^ (**Fig. 1a**). Certain bacterial species from the gut microbiome metabolize the bile acid pool exhaustively, mainly by hydrolyzing the taurine or glycine substituent on their sidechain (deconjugation) and then removing or epimerizing the hydroxyl groups at the 3 and 7 positions, producing dozens of structurally distinct secondary bile acids^3–11^. Some of these transformations are carried out by individual species, while others are ‘collaborative’ pathways between species. Roughly 95% of the bile acids released into the intestine are reabsorbed in the distal ileum and returned to the liver^12,13^, where some microbial modifications persist and others are enzymatically reversed. Bile acids transit this enterohepatic cycle ∼10 times per day^14^.

**Figure 1:**
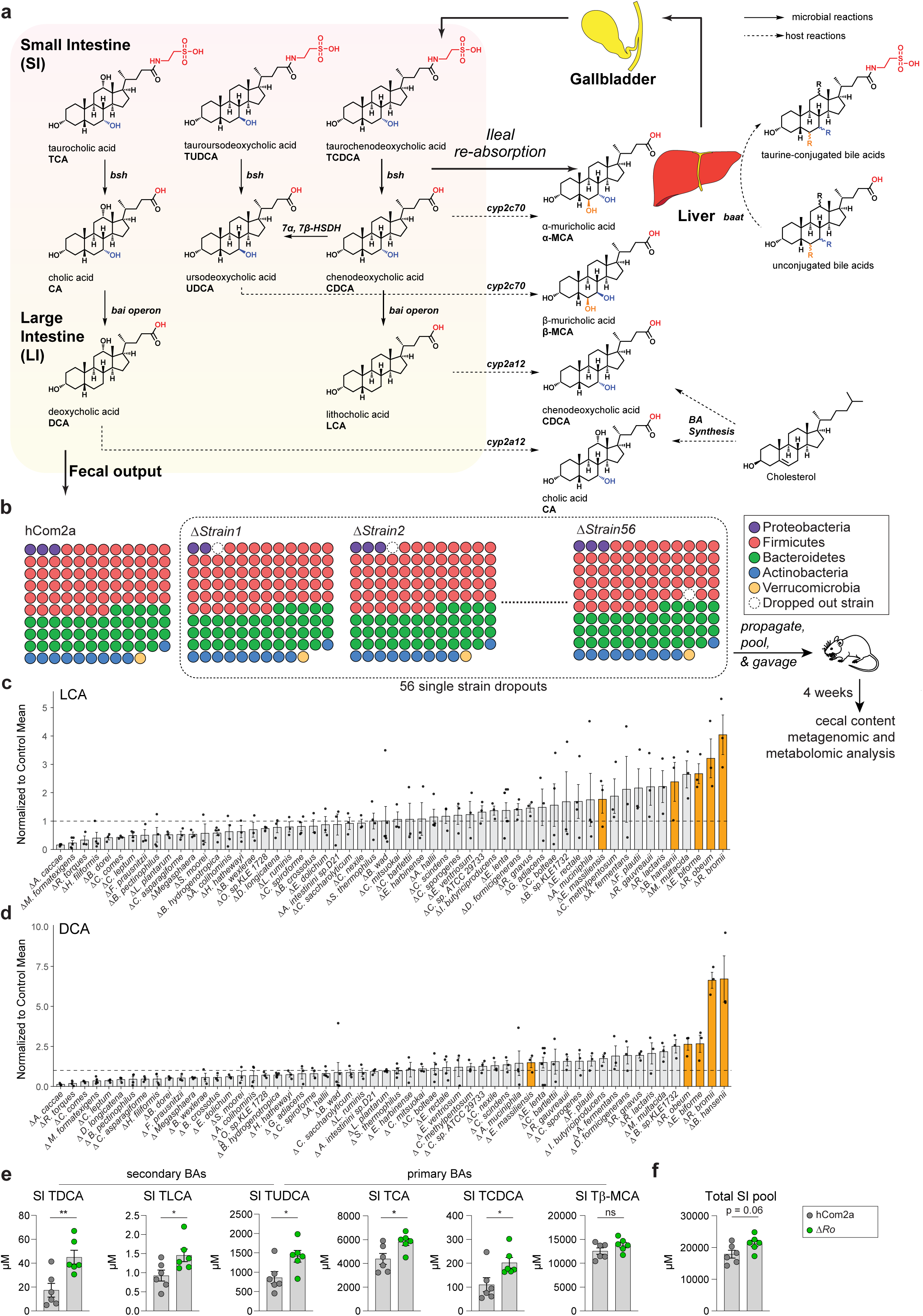
A single-strain dropout screen reveals communities that confer a larger bile acid pool on mice. (**a**) Schematic of the bile acid pool in the context of enterohepatic circulation. In the intestine, taurine-conjugated bile acids are deconjugated by microbial bile salt hydrolases, generating CA, CDCA, and UDCA. CA and CDCA undergo 7α-dehydroxylation by bacteria harboring the *bai* operon to produce the secondary bile acids DCA and LCA, respectively. Alternatively, CDCA can be epimerized to UDCA by 7α-and 7β-hydroxysteroid dehydrogenases (HSDHs). A small fraction of bile acids is secreted in the feces, while the majority (∼95%) reabsorbed in the ileum and returned to the liver. In hepatocytes, cytochromes P450 further remodel the bile acid pool: CYP2A12 converts DCA and LCA back to CA and CDCA, and CYP2C70 generates α- and β-muricholic acids from CDCA or UDCA. In addition, the host re-conjugates unconjugated bile acids via BAAT and synthesizes primary bile acids (CA and CDCA) *de novo* from cholesterol. Processed bile acids are secreted into the gallbladder and released into the small intestine upon feeding, completing the enterohepatic cycle. (**b**) Schematic overview of the single-strain dropout screen. 56 single-strain dropout communities were constructed and used to colonize germ-free Swiss-Webster mice. After 4 weeks, cecal contents were harvested for bile acid profiling and metagenomic sequencing. Typically, 6-12 single dropout communities were constructed in each batch. (**c-d**) Waterfall plots depicting the LCA and DCA levels in each dropout community, normalized to the corresponding control group (n=3–6 per condition). Several single-strain dropout communities exhibit elevated levels of lithocholic acid (LCA) and deoxycholic acid (DCA) in the cecum. (**e**) Δ*Ro*-colonized mice have an increased bile acid pool size in the small intestine. Small intestinal contents were measured by targeted LC-MS. TUDCA is both a primary and secondary bile acid in mice as it is synthesized by the host and the microbiome (n=6 per group). (**f**) Total small intestinal bile acid pool size (representing the sum of all measured bile acid species) in mice colonized by hCom2a or Δ*Ro* (n = 6 per group). All graphs show mean +/- SEM. Statistical significance was determined using a two-tailed Student’s t-test; p < 0.05 (*), p < 0.01 (**).

These metabolites have a broad range of biological activities that fall into two categories. First, their biophysical properties—principally facial amphipathicity—enable them to solubilize lipids in the intestinal lumen and facilitate their absorption^13,15^. Second, they are ligands for a variety of nuclear receptors and GPCRs, through which they modulate various aspects of host metabolism and immunity^16–18^.

Dysregulation of the bile acid pool is implicated in metabolic syndrome, primary sclerosing cholangitis^19,20^, primary biliary cholangitis^21^, and metabolic dysfunction-associated steatohepatitis^22,23^. Bile acid irregularities are also thought to play a pathologic role in irritable bowel syndrome with diarrhea (IBS-D)^24,25^ and some cases of inflammatory bowel disease^26^ and *Clostridium difficile* infection^27^. Current therapeutic strategies aim to reshape the pool size and composition in one of two ways: by augmenting it with hydrophilic bile acids like ursodeoxycholic acid (UDCA), or by inhibiting bile acid synthesis via FXR agonism or provision of FGF19 analogs^28–30^. However, these approaches carry significant drawbacks including limited efficacy, hepatotoxicity, and cardiovascular risk.

One of the most interesting opportunities in microbiome metabolism is to alter the size and composition of the bile acid pool *in situ*, by changing microbial biochemistry. Efforts to remove or mutate 7α-dehydroxylating strains have shown that the bile acid pool can be manipulated rationally^31,32^, but communities bereft of secondary bile acids are likely pathological^24,25^. No study to date has modified the bile acid pool in a way that would be durable and potentially beneficial.

Here, in the process of screening single-strain dropout variants of a defined gut microbial community, we discovered unexpectedly that a subset of dropout communities exhibited increased levels of lithocholic acid. We traced this effect to the bile salt hydrolase activity of a single strain—*Lactobacillus plantarum* WCFS1—which was sufficient to expand the size of the bile acid pool. By introducing engineered strains of *Lactobacillus* into the community that express bile acid metabolic genes, we manipulated the bile acid pool in one of two ways: enlarging its size or diverting it toward the production of 7β (urso) bile acids. These results demonstrate that specific strains and genes from the gut microbiome can be manipulated rationally to reshape the bile acid pool *in situ*, which may have important implications for physiology in health and disease.

### Single-strain dropout communities with increased secondary bile acids

To characterize the contributions of individual strains to community metabolism and host physiology, we performed a dropout screen in which we omitted, one strain at a time, 56 strains from a defined community, hCom2a (**Fig. 1b**) [Zeng *et al.* 2026a]. We colonized germ-free Swiss-Webster mice with each of these communities; after four weeks, we sacrificed the mice, harvested cecal contents and urine, and subjected the samples to targeted metabolite profiling and high-resolution metagenomic analysis. To account for batch effects, bile acid levels in each dropout were normalized to those of the corresponding control (**fig. s1d, Table S3**).

There were two striking bile-acid-related results. The first was a selective increase in 7β (urso) bile acids in mice deficient in *Clostridium sp. ATCC 29733* (Δ*CspA*) (**fig. s1a, s1e-f**). We discuss this finding further in the **Supplementary Text**.

The second was a set of five dropout communities that shared a common phenotype: a substantial increase in the level of the 7α-dehydroxylation products deoxycholic acid (DCA) and lithocholic acid (LCA) (**Fig. 1c-d**); their substrates, cholic acid (CA) and chenodeoxycholic acid (CDCA), were unaffected (**fig. s1b-c**). Thus, a subset of communities exhibited a specific increase in secondary (i.e., microbially generated) bile acids in the cecum.

Since the screen was designed to be general [Zeng *et al.* 2026a], metabolites were only profiled in the cecum and small intestine. However, because the bile acid pool is transformed by microbes in the intestine and host enzymes in the liver, its composition varies depending on where it is measured. Since bile acid reabsorption takes place in the terminal ileum, the small intestine is a better place to measure the pool: it is more concentrated, and its composition is a better representation of the set of molecules that circulate enterohepatically.

We started by profiling bile acids in the small intestine of mice colonized by one of the dropouts, Δ*Ro* (*Ruminococcus obeum*), focusing on taurine conjugates, which predominate in the SI. As we had observed in the cecum, the levels of the taurine conjugates of DCA (TDCA) and LCA (TLCA) were higher. However, so too were the levels of TUDCA, TCA, and TCDCA, indicating a more general increase in bile acid levels. Indeed, the total size of the bile acid pool trended larger by a substantial amount (3.5 mM), consistent with a model in which the secondary bile acid phenotype we initially observed in the cecum of Δ*Ro* was actually an augmentation of the circulating bile acid pool (**Fig. 1e-f, fig. s2a-b**).

### Dropout communities with higher secondary bile acid levels exhibit an increase in *Lactobacillus plantarum*

Next, we sought to understand the microbial basis of the pool size phenotype. We started by analyzing high-resolution metagenomic sequence data from mice colonized with the dropout communities. None of the communities show a difference in the level of either of the strains in hCom2a that carry out 7α-dehydroxylation, *Clostridium scindens* and *Clostridium hylemonae,* ruling out a direct effect resulting from a change in the relative abundance of the major secondary bile acid producers. However, we observed that in four of the five communities, the relative abundance of *Lactobacillus plantarum* increased substantially, revealing a commonality among communities that confer the phenotype (**Fig. 2a-d**).

**Figure 2:**
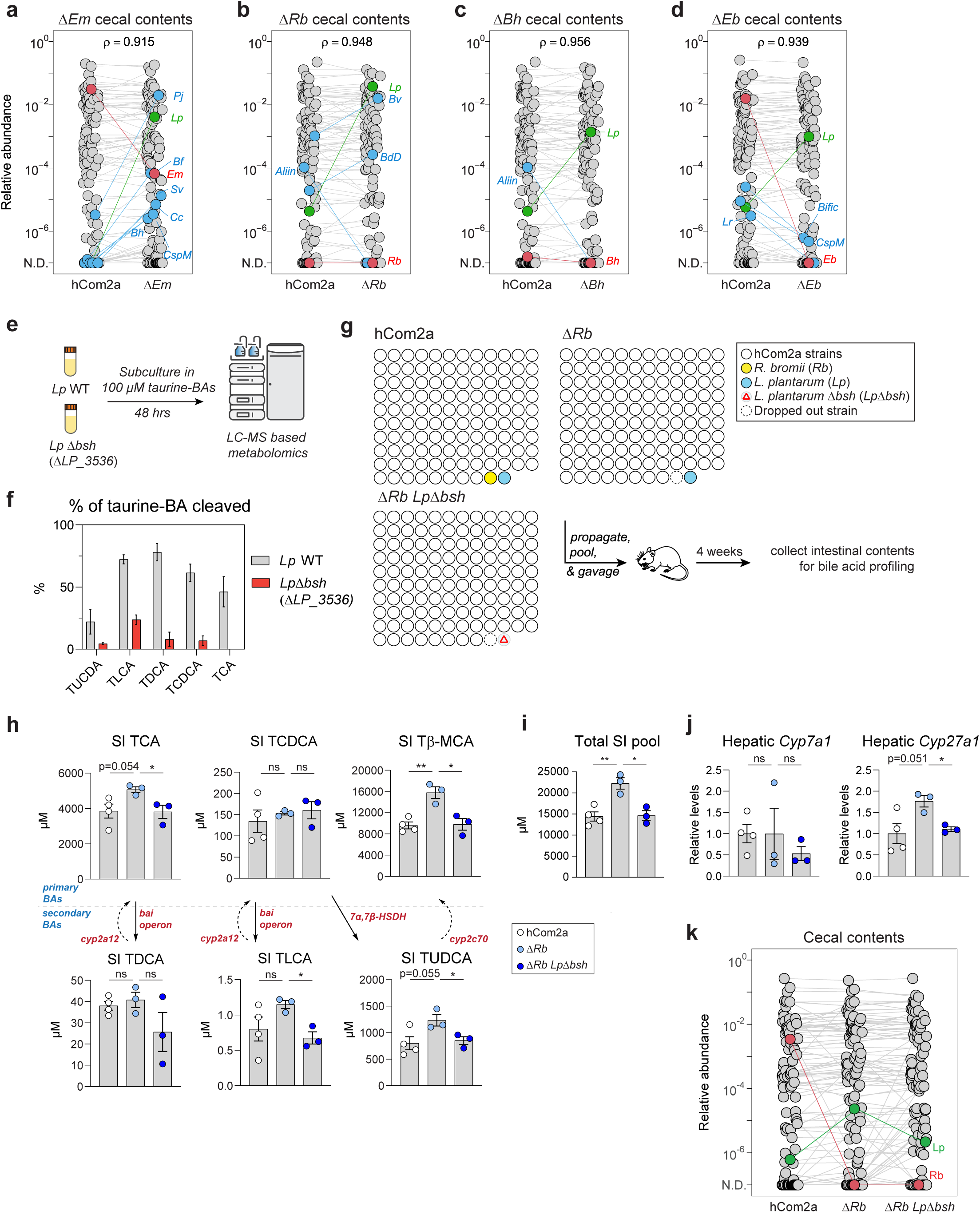
*Lactobacillus* bile salt hydrolase is responsible for the increased bile acid pool size. (**a-d**) Median relative abundances of hCom2a and Δ*Em* in cecal contents. Each dot is an individual strain; the collection of dots in a column represents the community with median values selected from 3-4 mice co-housed in one cage. The strain dropped out is highlighted in red; strains whose relative abundances change significantly are highlighted in blue (student t test p<0.05, fold change >10), and *Lactobacillus plantarum* (*Lp*) is highlighted in green. *Eisenbergiella massiliensis* (Δ*Em*)*, Ruminococcus bromii* (Δ*Rb*)*, Blautia hansenii* (Δ*Bh*), and *Eubacterium biforme* (Δ*Eb*) dropouts are shown. (**e**) Schematic of the experiment: *L. plantarum* WCFS1 and a *bsh*-deficient mutant (Δ*LP_3536*) were incubated with taurine-conjugated bile acids for 48 h. Bile acids were extracted from the media and analyzed by LC-MS to assess each strain’s ability to deconjugate bile acids. (**f**) LP_3536 is the major bile salt hydrolase in *L. plantarum* WCFS1. The percentage of each bile acid species being deconjugated after incubation with *L. plantarum* WCFS1 (WT) or Δ*LP_3536* for 48 h is shown. (**g**) Schematic of *in vivo* experiments testing the role of *L. plantarum bsh*. Two variants of hCom2a were constructed: one in which *Rb* is dropped out (Δ*Rb*), and a second in which *Rb* is omitted and wild-type *L. plantarum* is replaced with a *bsh*-deficient mutant (Δ*Rb Lp*Δ*bsh*). hCom2a, Δ*Rb,* and Δ*Rb Lp*Δ*bsh* were used to colonize germ-free Swiss-Webster mice. After four weeks, intestinal contents were harvested for bile acid profiling and metagenomic sequencing. (**h-i**) Targeted metabolomic analysis of small intestinal contents. In Δ*Rb* there is a general increase in bile acid levels that leads to an increase in the size of the bile acid pool; this effect is reversed in the absence of *L. plantarum bsh* (n=3-4 per group). (**j**) Gene expression analysis for *Cyp7a1* and *Cyp27a1* in the liver (n=3-4 per group). (**k**) Median relative abundances of hCom2a, Δ*Rb*, and Δ*Rb Lp*Δ*bsh* in the cecal contents of colonized mice. Each dot is an individual strain; the collection of dots in a column represents the community with median values selected from 3-4 mice co-housed in one cage. All graphs show mean +/- SEM. Statistical significance was determined using a two-tailed student’s t-test; p < 0.05 (*), p < 0.01 (**).

### *Lactobacillus plantarum* bile salt hydrolase is responsible for the increase in secondary bile acids

*L. plantarum* is known to carry out a different bile acid transformation that is upstream of 7α-dehydroxylation: hydrolysis of the host-derived primary bile acids tauro-CA (TCA), glyco-CA (GCA), tauro-CDCA (TCDCA), and glyco-CDCA (GCDCA) to generate their free carboxylic acid derivatives, CA and CDCA (**Fig. 1a**). We hypothesized that the enzyme that carries out this transformation—bile salt hydrolase (*bsh*)—plays a role in the bile acid pool size phenotype we had observed. To test this hypothesis, we started by constructing a mutant of *L. plantarum* deficient in bile salt hydrolase activity. The strain of *L. plantarum* in our community, *L. plantarum* WCFS1, harbors four putative bile salt hydrolase genes; a mutant deficient in one of them, LP_3536, is known to have a reduced capacity to deconjugate taurine-conjugated bile acids^33^. We recreated this mutant—here termed *Lp*Δ*bsh*—and confirmed that it is largely (but not completely) deficient in the ability to deconjugate TUDCA, TLCA, TDCA, TCDCA, and TCA (**Fig. 2e-f**).

Next, we constructed two sets of two communities. The first was one of the single-strain dropouts in which we observed the phenotype of increased secondary bile acid levels (Δ*Rb*), and the second was a derivative of Δ*Rb* in which we replaced *L. plantarum* with the Δ*bsh* mutant (Δ*Rb Lp*Δ*bsh*). Thus, the second community differs from hCom2a by the absence of one strain (*Ruminococcus bromii, Rb*) and one gene (*bsh*) from a second strain (*Lp*) (**Fig. 2g**).

We colonized germ-free Swiss-Webster mice with hCom2a, Δ*Rb*, and Δ*Rb Lp*Δ*bsh*, waited four weeks, and then measured bile acid levels in the small intestine. In Δ*Rb* (as observed previously in Δ*Ro*), there is a general increase in bile acid levels that leads to an increase in the size of the bile acid pool. Notably, this phenotype is reversed by the removal *L. plantarum bsh* (**Fig. 2h-i**); the levels of TCA, Tβ-MCA, TLCA, and TUDCA in Δ*Rb Lp*Δ*bsh* are 1 mM, 6 mM, 0.5 mM, and 0.4 mM lower than in Δ*Rb*. Overall, eliminating the *bsh* gene from *Lp* led to a 7 mM reduction in the size of the bile acid pool.

On the one hand, this is consistent with our hypothesis that bile salt hydrolase plays a role in the phenotype of increased secondary bile acids. On the other hand, the direction of the effect was not immediately obvious: one might have anticipated that the removal of bile salt hydrolase would increase (not decrease) the level of its substrates, conjugated bile acids. To determine why deleting bile salt hydrolase from *L. plantarum* led to a decrease in conjugated bile acids, we measured the level of hepatic *Cyp7a1* and *Cyp27a1*, the rate-limiting enzymes for the classical and alternative bile acid synthesis pathways. In mice colonized by Δ*Rb*, *Cyp27a1* was induced; this induction was reversed when *bsh* was deleted from *L. plantarum* (**Fig. 2j**). The same pattern was not observed for *Cyp7a1*. Thus, while *L. plantarum* bile salt hydrolase cleaves taurine-conjugated bile acids, it also triggers an increase in bile acid synthesis via the alternative pathway, leading to an increase in conjugated bile acids.

To assess how the strain dropout and gene deletion communities compare to hCom2a in terms of strain relative abundances *in vivo*, we performed a high-resolution metagenomic analysis of the cecal contents from these mice. As observed in the initial screen, the relative abundance of *Lp* increases when *Rb* is omitted. Notably, in Δ*Rb Lp*Δ*bsh*, the relative abundance of the *Lp* Δ*bsh* mutant is substantially lower than in the corresponding community with wild-type *Lp* (p = 0.017, Δ*Rb vs* Δ*Rb Lp*Δ*bsh*) (**Fig. 2k**). These results suggest that bile salt hydrolase activity confers a competitive advantage on *L. plantarum* in the context of a complex community *in vivo*.

### Controlling bile acid pool size by community-scale *bsh* engineering

Having observed that deleting *bsh* from *L. plantarum* can reduce the size of the bile acid pool, we sought to determine whether a similar maneuver could generate the opposite effect: if we add more bile salt hydrolase activity to hCom2a, can we increase the pool size beyond the point observed in our single-strain dropout communities?

We reasoned that the simplest way to increase the bile salt hydrolase activity of hCom2a would be to add additional strains of *Lactobacillus* that harbor *bsh*. To test this hypothesis, we screened a panel of *bsh*-positive *Lactobacillus* strains in vitro for their ability to hydrolyze taurine-conjugated bile acids (TUDCA, TLCA, TDCA, TCDCA and TCA), releasing taurine and a deconjugated bile acid (**Fig. 3a**). All of the strains we tested carried out bile salt hydrolysis, with *L. plantarum* WCFS1 (already in hCom2a), *L. plantarum* MAF1421, *L. gasseri* ATCC 33323, *L. acidophilus* ATCC 4356, and *L. reuteri* I49 exhibiting the strongest activity (**Fig. 3b**). Based on these results, we constructed two variants of hCom2a: one in which we added the four *Lactobacillus* strains not already present in hCom2a (hCom2a + L4A), and another in which we deleted the resident strain of *Lactobacillus plantarum* strain (Δ*Lp*) (**Fig. 3c**). We colonized one group of germ-free SW mice with this community and another with Δ*Lp* as a control. After four weeks, we profiled bile acid levels in the cecum and small intestine.

**Figure 3:**
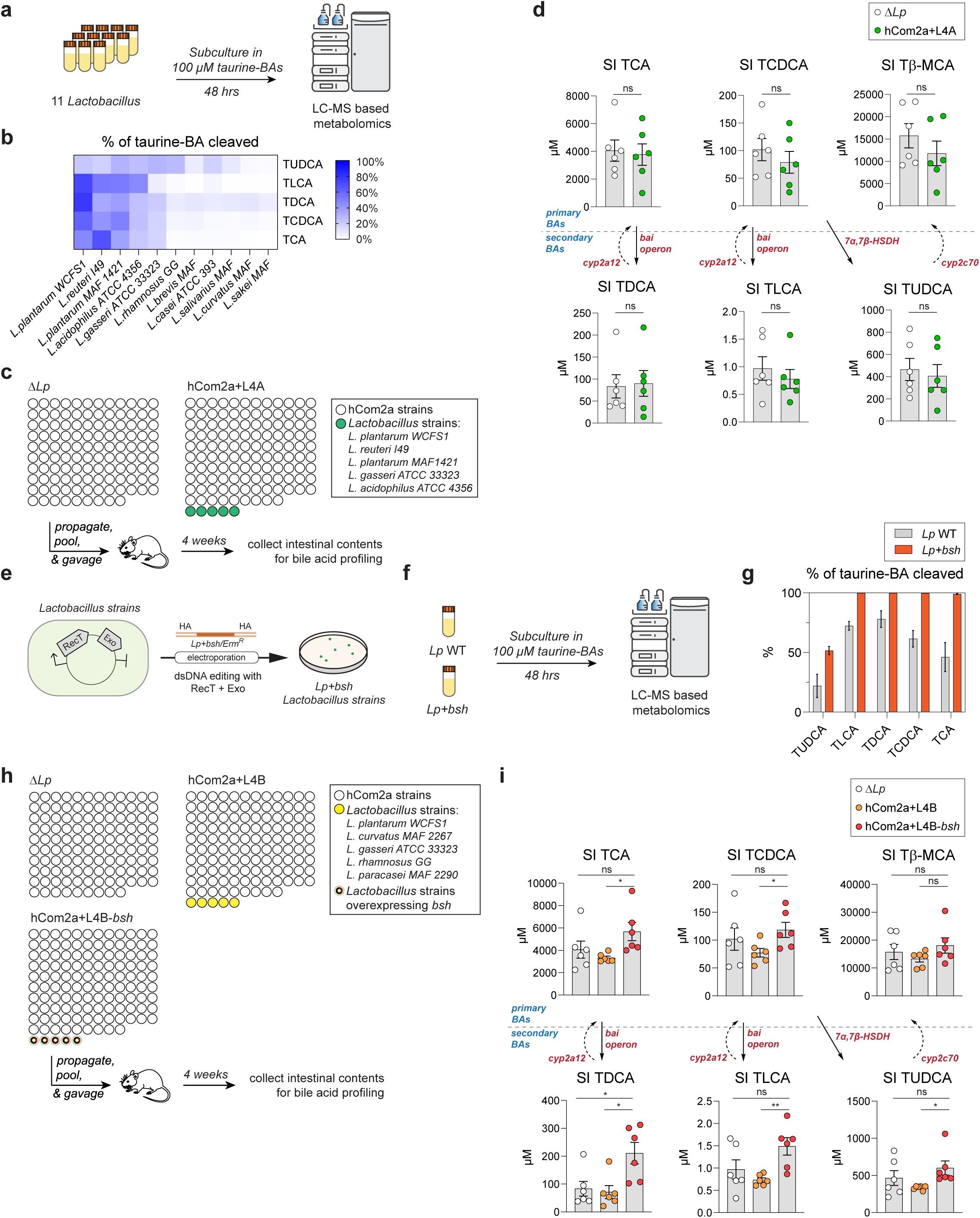
Community-level *bsh* engineering increases secondary bile acid production. (**a**) Schematic of the experiment. Eleven *Lactobacillus* strains were incubated with taurine-conjugated bile acids for 48 h; the extent of deconjugation by each strain was assessed by targeted LC-MS. (**b**) Bile acid deconjugation by a panel of *Lactobacillus* species. The extent of deconjugation differed by strain and by bile acid species. (**c**) Schematic of an experiment to test whether augmenting hCom2 with additional *bsh-*expressing *Lactobacillus* species increases secondary bile acid levels in mice. Two derivatives of hCom2a were constructed: one in which *L. plantarum* WCFS1 was removed (Δ*Lp*), and another in which four *bsh-*expressing *Lactobacillus* species were added, for a total of five (hCom2a+L4A). Germ-free mice were colonized for 4 weeks; mice were then sacrificed and cecal contents and small intestinal contents were then subjected to metabolomic and metagenomic analysis. (**d**) Targeted metabolomic analysis of small intestinal contents. The addition *bsh*-expressing *Lactobacillus* (hCom2a+L4A) did not increase secondary bile acid levels compared to Δ*Lp* (n=6 per group). (**e**) Five *Lactobacillus* strains were engineered to overexpress *bsh* using RecT/RecE-mediated recombineering. (**f**) Wild-type *L. plantarum* WCFS1 and a mutant overexpressing *bsh* (*Lp*+*bsh*) were incubated with taurine-conjugated bile acids for 72 h. Bile acids were extracted from the medium and analyzed by LC-MS to assess each strain’s ability to deconjugate bile acids. (**g**) *Lp*+*bsh* deconjugated bile acids more efficiently than wild-type *L. plantarum* WCFS1. The percentage of each bile acid species converted after incubation with *Lp* WT *or Lp+bsh* for 48 h is shown. (**h**) Three variants of hCom2a were constructed: one in which *L. plantarum* was omitted (Δ*Lp*), a second in which *L. plantarum* and four additional species of *Lactobacillus* were added (hCom2a+L4B), and a third that is identical to the second except each strain of *Lactobacillus* was engineered to overexpress *bsh* (hCom2a+L4B-*bsh*). Germ-free mice were colonized with each community for 4 weeks; mice were then sacrificed and cecal contents and small intestinal contents were subjected to metabolomic and metagenomic analysis. (**i**) Targeted metabolomics of small intestinal contents. Mice colonized with hCom2a+L4B-*bsh* had higher levels of secondary bile acid levels than both control groups (n=6 per group). All graphs show mean +/- SEM. Statistical significance was determined using a two-tailed Student’s t-test; p < 0.05 (*), p < 0.01(*).

In both sites, we saw no difference in the level of secondary bile acid species (**Fig. 3d, fig. s3a**). No clear microbiologic signal helped to explain this result. In mice colonized by the hCom2a+ L4A community, *L. reuteri I49* had a high relative abundance, while the four other *Lactobacillus* strains remained below the detection limit in the cecum (**fig. s3b-c**). Since *Lactobacillus* species preferentially colonize the small intestine, their abundance may be underestimated when sampling from the cecum (**fig. s3g-h**). Nevertheless, these data suggest that augmenting the community with more *bsh*-positive *Lactobacillus* strains is not, on its own, sufficient to impact bile acid metabolism.

We speculated that this may reflect a fixed niche size for *Lactobacillus*, limiting the total carrying capacity. If so, by adding more *Lactobacillus* species, we would not be able to increase the net amount of bile salt hydrolase activity in the intestine. However, we hypothesized that we could still increase total BSH activity by making each strain more efficient at carrying out this reaction.

To test this hypothesis, we engineered five strains of *Lactobacillus* to express *bsh* under the control of the strong promoter P20787^34^ (**Fig. 3e**); the engineered strains exhibit a stronger bile salt hydrolase activity than their parental WT counterparts (**Fig. 3f-g**). We then constructed two variants of hCom2a: one that includes these engineered strains (hCom2a+L4B-*bsh*) and another that includes the wild-type versions of the same five strains, without the transgene (hCom2a+L4B). We colonized two groups of germ-free SW mice with these communities (and another with Δ*Lp* as a control), waited four weeks, and profiled bile acids in the cecum and small intestine (**Fig. 3h, fig. s3e-f**). In the cecum, we saw a specific increase in the level of secondary bile acids (DCA, LCA, and isoDCA) but not primary bile acids, similar to the original observation from the strain dropout experiment (**fig. s3d**). In the small intestine, there was a marked increase in TDCA and TLCA, with the former up almost 3-fold (an increase of >100 µM) (**Fig. 3i**). These data demonstrate that augmenting bile salt hydrolase activity—through the addition of a single transgene in five strains in the community—can lead to a substantial increase in the bile acid output of the community.

### Characterizing microbial and host effects on the bile acid pool

Our efforts to model human bile acid metabolism in mice are impeded by two transformations that are specific to mice and skew the bile acid pool substantially from its composition in humans. The first is the 6-hydroxylation of CDCA and UDCA by CYP2C70 to form α- and β-muricholic acid, both of which are absent in humans and the latter of which is routinely the most abundant bile acid in the murine pool^35^. The second is the 7α-hydroxylation of DCA and LCA by CYP2A12 to regenerate CDCA and CA, reversing 7α-dehydroxylation and skewing the pool toward primary bile acids^36^. These differences limit our ability to study microbial bile acid metabolism—especially the production of LCA, UDCA, and their derivatives—and model the effects of engineering the microbiome on the bile acid pool.

To overcome these limitations, we used adeno-associated viral CRISPR (AAV-CRISPR) to disrupt the murine *Cyp2c70* and *Cyp2a12* genes^15^ (**Fig. 4a**). Intraperitoneal injections of an AAV encoding Cas9 and a guide RNA led to gene mutation and loss of protein function in the liver within one week, making this an ideal tool for gene disruption in gnotobiotic mice colonized by defined communities. AAV-CRISPR constructs that target each of these genes were recently developed and validated in mice^15^.

**Figure 4:**
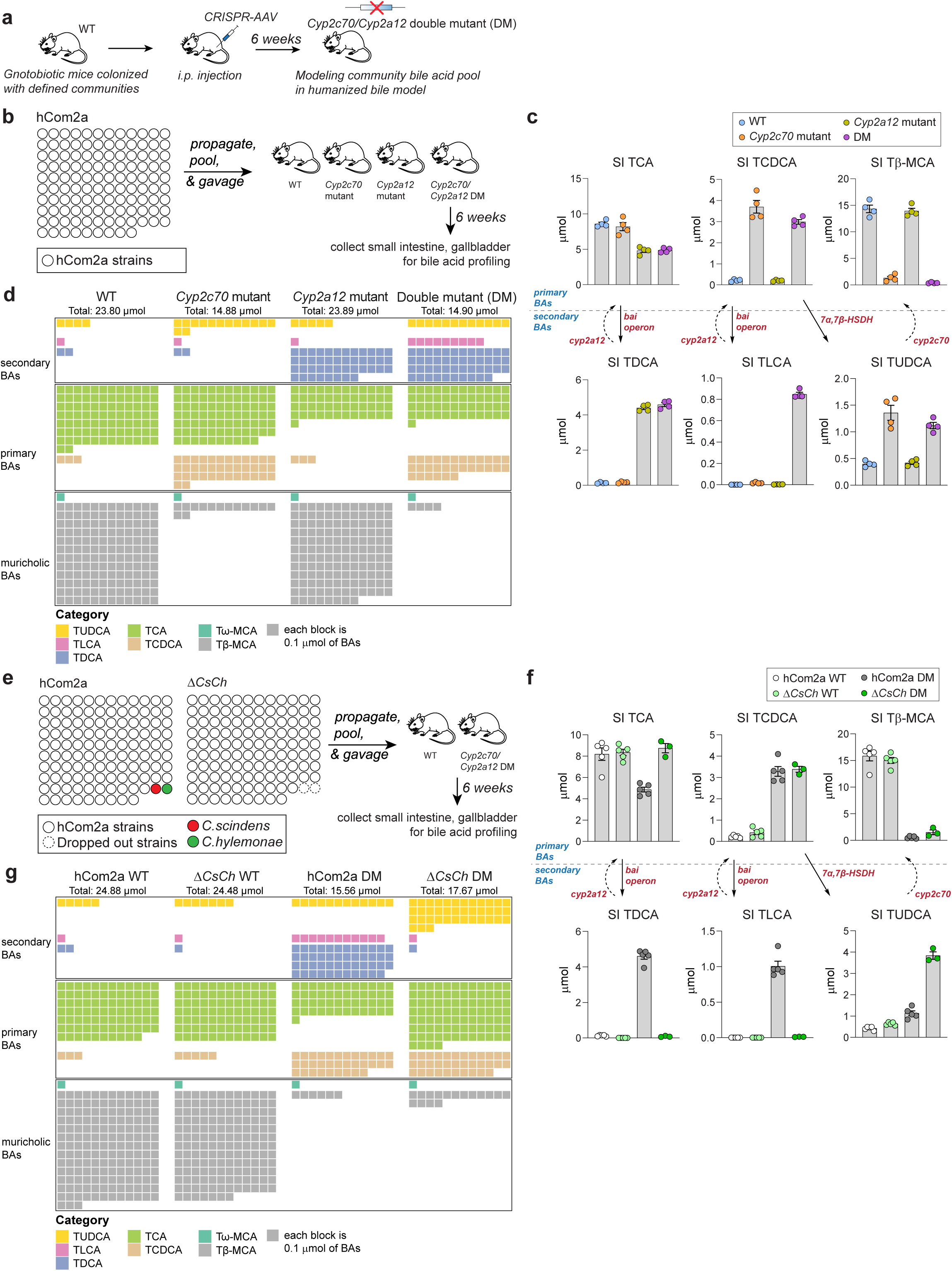
Characterizing microbial and host effects on the bile acid pool. (**a**) Gnotobiotic mice colonized by defined communities were injected with AAV-CRISPR lines targeting two host genes, *Cyp2c70* and *Cyp2a12*, in order to ‘humanize’ the bile acid pool by eliminating murine-specific transformations. (**b**) Four groups of mice were colonized hCom2a; they were injected i.p. with engineered AAVs targeting *Cyp2a12*, *Cyp2c70*, a combination of the two, or control AAVs. Mice were housed for 6 weeks, sacrificed, and samples were collected and subjected to metabolomic and metagenomic analysis. (**c**) Targeted metabolomic analysis of small intestinal contents. The bile acid pool in hCom2a-colonized mice is altered dramatically by mutating *Cyp2a12*, *Cyp2c70,* or both, and in the double mutant it more closely resembles the human bile acid pool (n = 4 per group). (**d**) A waffle plot showing a visual representation of the bile acid pool in the mice from the experiment schematized in (**b**). Each block corresponds to 0.1 μmol of bile acids. (**e**) Two communities were constructed: hCom2a and a double-dropout derivative in which the two strains capable of carrying out 7α-dehydroxylation, *Clostridium scindens* and *Clostridium hylemonae*, were removed (Δ*CsCh*). Each community was used to colonize two groups of germ-free mice: one was injected *i.p.* with control AAVs, and the other engineered AAVs targeting *Cyp2a12* and *Cyp2c70* (double mutant, DM). Mice were housed for 6 weeks and then sacrificed; samples were collected and subjected to metabolomic and metagenomic analysis. (**f**) Targeted metabolomic analysis of small intestinal contents. The removal of *Cs* and *Ch* resulted in a marked reduction of TDCA and TLCA and a compensatory increase in TCDCA and TUDCA, an effect that is only evident in mice with a humanized bile acid pool (n=5 per group for WT, n=4 for hCom2a DM, and n=3 for Δ*CsCh* DM). (**g**) A waffle plot showing a visual representation of the bile acid pool in the mice from the experiment schematized in (**e**). Each block corresponds to 0.1 μmol of bile acids. All graphs show mean +/- SEM.

Germ-free mice colonized with hCom2a were injected with AAV-CRISPR constructs that target *Cyp2c70*, *Cyp2a12*, or a mixture of both. After waiting six weeks for muricholic acids to be cleared from enterohepatic circulation, we analyzed bile acid profiles from the small intestine of each group of mice (**Fig. 4b**). We made three major observations. First, *Cyp2c70* disruption largely eliminates muricholic acid production and leads to compensatory increases in TUDCA and TCDCA, consistent with prior studies of germline mutants^15,17,36,37^. The overall pool size was smaller in the absence of CYP2C70, likely due to the loss of muricholic acid, which normally derepresses hepatic bile acid synthesis by antagonizing FXR. Second, *Cyp2a12* mutant mice have increased TDCA and reduced TCA levels, consistent with the role of CYP2A12 in reversing 7α-dehydroxylation^17,37^; other bile acid levels remain largely unchanged. Third, the simultaneous mutation of *Cyp2c70* and *Cyp2a12* (DM) yields a bile acid pool with both properties: reduced muricholic acid levels and increased secondary bile acids, more closely resembling the bile acid pool of a typical human. Notably, in the DM mice (but not in either of the single mutants), TLCA levels increase significantly (∼1 µmol), suggesting that lithocholic acid accumulation requires the absence of both murine-specific P450 enzymes. This is likely because LCA can be reverted to CDCA by CYP2A12, and CYP2C70-mediated conversion of CDCA to UDCA competes with 7α-dehydroxylation for the formation of LCA (**Fig. 4c-d, fig. s4a-b**).

Overall, community composition was consistent across murine genetic backgrounds, with only a few exceptions: *B. uniformis* increased in relative abundance in the *Cyp2c70* mutant, while *D. formicigenerans* was less abundant in the double mutant background (**fig. s4c-d**). These observations suggest that most strains colonize independently of bile acid composition, though a few are sensitive to changes in the composition of the bile acid pool.

DCA and LCA, both products of 7α-dehydroxylation, are the most abundant microbiome-derived bile acids and constitute a large portion of the human bile acid pool^4^. In previous work, we constructed a community lacking two strains that carry out 7α-dehydroxylation, *Clostridium scindens* (*Cs*) and *Clostridium hylemonae* (*Ch*), and observed a reduction in 7α-dehydroxylation products^31^. However, the effect of modulating microbial bile acid production has not been characterized in the setting of a humanized bile acid pool. To start, we constructed hCom2a and a double dropout community lacking *Cs* and *Ch* (Δ*Cs*Δ*Ch*) and used them to colonize two groups of germ-free mice each. One group of mice in each colonization state was not modified further (hCom2a WT and Δ*Cs*Δ*Ch* WT), while the other was subjected to AAV-CRISPR mediated inactivation of *Cyp2a12* and *Cyp2c70* (hCom2a DM and Δ*Cs*Δ*Ch* DM) (**Fig. 4e, fig. s4g-h**).

In WT and *Cyp2a12*/*Cyp2c70* double mutant mice, removal of *Cs* and *Ch* resulted in a marked diminution of TDCA, confirming the loss of 7α-dehydroxylation activity. In double mutant mice colonized by Δ*Cs/Ch*, TLCA was largely eliminated as well, leading to a compensatory increase in TCDCA and TUDCA. The bile acid pool in the double mutant mice was profoundly affected by the absence of *Cs* and *Ch*: ∼36% of the pool (TDCA and TLCA) was eliminated, with corresponding increases in TCA and TUDCA. These shifts were far less pronounced in WT mice (**Fig. 4f-g, fig. s4e-f**). Thus, eliminating microbial 7α-dehydroxylation leads to a substantial redistribution of the bile acid pool, an effect that is only evident in mice with a humanized bile acid pool.

### Engineering the pool toward urso bile acids

Cholestatic disease is caused by an obstruction of bile flow from the liver to the small intestine. UDCA is an FDA-approved treatment; it is thought to work by rendering the bile acid pool more hydrophilic and decreasing its detergent-like properties. However, the provision of exogenous UDCA enlarges the pool, which could be counterproductive in the setting of cholestasis. We hypothesized that it might be more effective to engineer the microbiome to divert the bile acid pool toward UDCA *in situ*, rendering it more hydrophilic without adding exogenous bile acids^38^.

Inspired by our observation of increased UDCA in the *Clostridium sp.* ATCC 29733 (Δ*CspA*) dropout community (**fig. s1e-i**, **Supplementary Text**), we sought to selectively increase UDCA by rational means. Two considerations guided our approach. First, *R. gnavus*—the strain most likely responsible for increased UDCA in Δ*CspA* (**fig. s1e-i**, **Supplementary Text**)—is not genetically tractable. Second, the increased UDCA in Δ*CspA*-colonized mice was primarily observed in the cecum, outside of enterohepatic circulation. We reasoned that a better strategy might be to position 7α- and 7β-HSDH enzymes in the small intestine, enabling the substrate (CDCA) to be converted to UDCA before being consumed by competing pathways, primarily 7α-dehydroxylation.

To do so, we needed a microbial host in which to express these enzymes. After considering various options, we chose *Lactobacillus*: its species preferentially colonize the small intestine and they have sufficient metabolic ‘leverage’ to carry out this pathway, having served as the host for our efforts to modulate pool size by manipulating bile salt hydrolase. We hypothesized that by overexpressing 7α- and 7β-HSDH in *Lactobacillus*, we could skew the pool toward urso bile acids.

We codon-optimized 7α- and 7β-HSDH genes from *R. gnavus* for *Lactobacillus* and placed them under the control of a strong promoter in an artificial operon (**Fig. 5a**). This construct was integrated into the genome of eight *Lactobacillus* strains. As expected, the edited strains readily convert CA to 7β-CA (ursocholic acid) and CDCA to UDCA *in vitro* (**Fig. 5b-c**). To test their activity *in vivo*, we added all eight edited strains to Δ*Lp*, yielding a community that differs by only two genes (in each of eight strains), hCom2a-L7-urso. We also constructed a control community, hCom2a-L7, in which the wild-type versions of the same eight *Lactobacillus* strains were added to Δ*Lp* (**Fig. 5d**).

**Figure 5:**
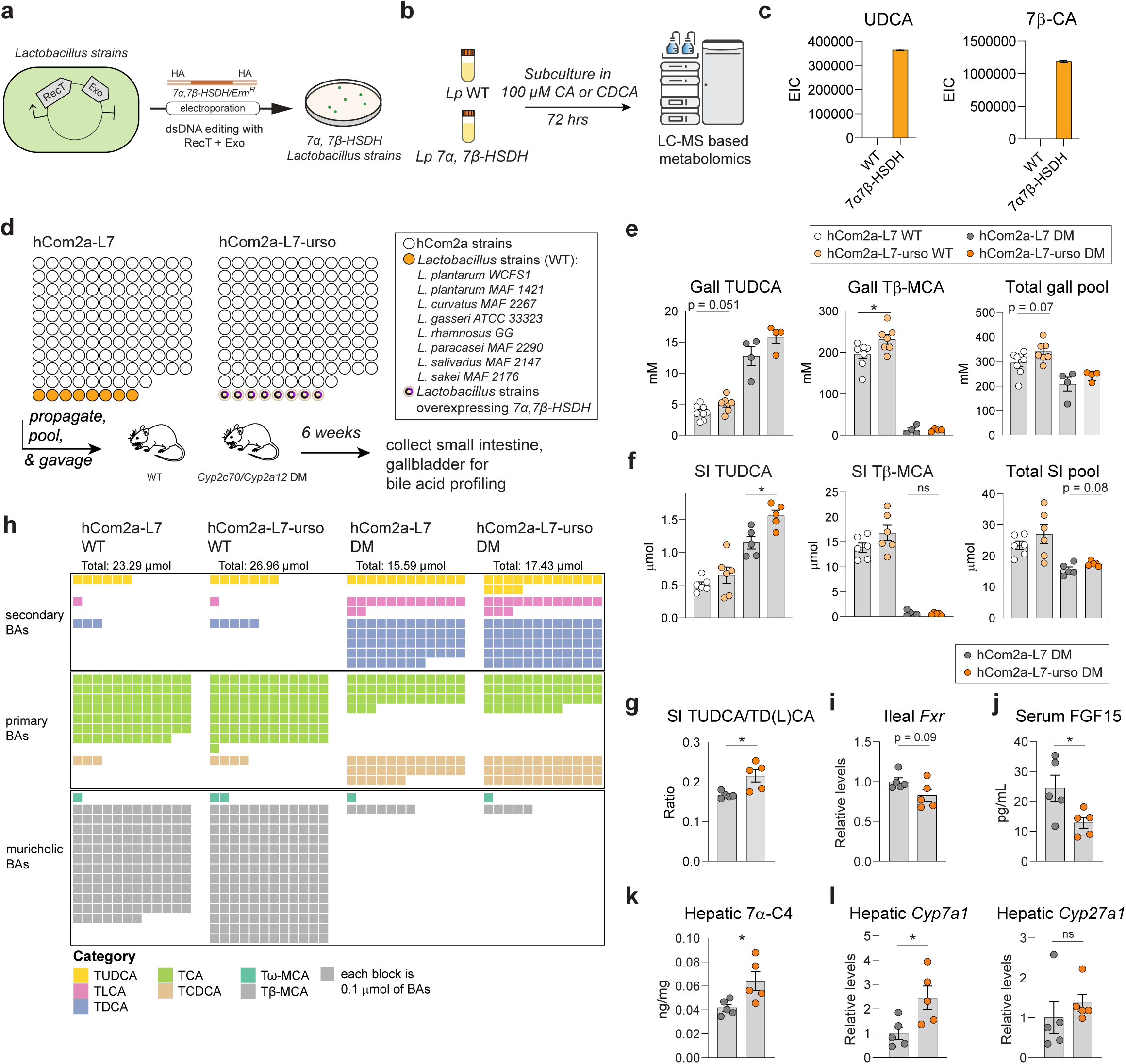
Engineering the pool toward urso bile acids. (**a**) Eight *Lactobacillus* strains were engineered to overexpress 7α- and 7β-HSDH using RecT/RecE-mediated recombineering. (**b**) Wild-type *L. plantarum* WCFS1 and a mutant overexpressing 7α- and 7β-HSDH were incubated with CA or CDCA for 72 h. Bile acids were extracted from the medium and analyzed by LC-MS to assess each strain’s ability to epimerize the 7-OH. (**c**) *Lp*+7αα,β-HDSH generated urso bile acids more efficiently than wild-type *L. plantarum* WCFS1. An extracted ion chromatogram showing the percentage of each bile acid species converted after incubation with *Lp* WT *or Lp*+7αα,β-HDSH for 72 h is displayed, n=2. (**d**) Two variants of hCom2a were constructed: one in which seven additional species of *Lactobacillus* were added for a total of eight (hCom2a-L7), and a second that is identical to the first except each strain of *Lactobacillus* was engineered to overexpress 7α- and 7β-HSDH (hCom2a-L7-urso). Each community was used to colonize two groups of germ-free mice: one was injected i.p. with control AAVs, and the other engineered AAVs targeting *Cyp2a12* and *Cyp2c70* (double mutant, DM). Mice were housed for 6 weeks, sacrificed, and cecal contents and small intestinal contents were subjected to metabolomic and metagenomic analysis. (**e**) Targeted metabolomics of gallbladder contents. Concentrations of TUDCA and Tβ-MCA and total gallbladder bile acid contents were quantified by targeted LC-MS based metabolomics. hCom2a-L7-urso had higher Tβ-MCA and a trend toward increased TUDCA and a larger bile acid pool (n=8 for hCom2a WT, n=7 for hCom2a-urso WT, n=4 for DM groups). (**f**) Targeted metabolomics of small intestinal contents. TUDCA, Tβ-MCA, and total bile acid pool in the small intestine were quantified via targeted LC-MS based metabolomics. hCom2a-L7-urso had higher UDCA and a trend toward an increased bile acid pool (n=6 for WT groups, n=5 for DM groups). (**g**) The ratio of TUDCA to the products of the competing 7α-dehydroxylation pathway (TDCA+TLCA) was increased in hCom2a-L7-urso-colonized mice (n=5). (**h**) A waffle plot showing a visual representation of the bile acid pool in the small intestine of mice from the experiment schematized in (**d**). Each block corresponds to 0.1 μmol of bile acids. (**i**) Gene expression analysis of ileal *Fxr* (n=5 per group). (**j**) Serum FGF15 concentration measured by ELISA (n=5 per group). (**k**) Hepatic 7α-C4 concentration measured by LC–MS (n=5 per group). (**l**) Gene expression analysis of hepatic *Cyp7a1* and *Cyp27a1* (n=5 per group). All graphs show mean ± SEM. Statistical significance was determined using a two-tailed Student’s t-test; p < 0.05 (*).

We colonized wild-type and *Cyp2a12*/*Cyp2c70* double mutant mice with hCom2a-L7 and hCom2a-L7-urso, waited 6 weeks, and profiled bile acids in the small intestine and gallbladder (**Fig. 5d**, **fig. s5d-e**). Wild-type mice colonized by hCom2a-L7-urso had a large (36 mM) increase in the 6-hydroxylation product of UDCA, β-muricholic acid, in the gallbladder (**Fig. 5e-h, fig. s5a-c**). These data very likely indicate increased production of UDCA, and they demonstrate that the abundance of a single bile acid species can be increased rationally.

In double mutant mice colonized by hCom2a-L7-urso, TUDCA levels increased modestly—from ∼1.0 µmol to 1.5 µmol, favoring 7α-epimerization over 7α-dehydroxylation (TDCA and TLCA) (**Fig. 5g**). To our surprise, TDCA, TCA, and TCDCA increased as well, by a similar amount. Thus, the addition of enzymes that generate TUDCA to the microbiome unexpectedly led to an increase in several bile acid species and an expansion (12% in the small intestine and 16% in the gallbladder) of the bile acid pool (**Fig. 5e-h, fig. s5a-c**).

We hypothesize that the accumulation of TUDCA antagonizes intestinal FXR. This would de-repress hepatic bile acid synthesis via reduced FGF15 signaling, supplying more substrates to downstream pathways. To test this hypothesis, we measured ileal *Fxr* mRNA, hepatic *Cyp7a1* and *Cyp27a1* mRNA, serum FGF15 protein, and hepatic 7α-hydroxy-4-cholesten-3-one (7α-C4) in double-mutant mice. We observed a trend toward decreased *Fxr* expression and circulating FGF15 protein levels, along with increased *Cyp7a1* expression and hepatic 7α-C4. (**Fig. 5i-l**), consistent with UDCA–mediated expansion of the bile acid pool through the FXR–FGF15–*Cyp7a1* axis. Together, these findings demonstrate that rationally designed communities can alter the composition of the bile acid pool, though sometimes in unexpected ways.

## DISCUSSION

Genetic screens often follow a simple logic: removing a gene eliminates its protein product, triggering a phenotype related to the loss of its function. The microbiologic screen that gave rise to this project differed in two important ways. First, although the screen involved removing strains, the phenotype was not a loss but a gain of function: a rise in secondary bile acid levels caused by increased bile salt hydrolase activity. Second, it came about indirectly; it was not caused by the absence of the omitted strains, but by a *Lactobacillus* that bloomed when they were removed. Thus, while strain dropouts are an informative tool to probe community-scale metabolism, their logic is often indirect—via an ecological shift—and their results can be hard to predict.

Nevertheless, the phenotypes were quite large: In our initial screen, the removal of individual strains triggered shifts of 3-6 mM in the small intestinal bile acid pool (**Fig. 1f**, **Fig. 2i**). Communities engineered to express *bsh* led to a 2–3-fold increase in TDCA and TLCA and a 6 mM increase in pool size (**Fig. 3i**), while humanized mice colonized by 7α/β-HSDH-expressing communities had a profound expansion of the bile acid pool—36 mM in the gallbladder (**Fig. 5e, h**). Moreover, unlike in previous work, including our own^31,32,39^, we brought these changes about without eliminating the 7α-dehydroxylation niche (which would likely be unstable) or introducing a pathobiont like *E. coli*^40^. The maneuvers we used—adding or removing individual strains and adding a transgene to a native colonist like *Lactobacillus*—would in principle be safe and would not leave any niches open, increasing the likelihood of stability if used therapeutically.

An ecological shift upon strain removal gave rise to the metabolic phenotype we studied here. At the same time, expression of bile salt hydrolase confers a growth advantage on *Lactobacillus* in the small intestine (**Fig. 2k**). Thus, the connection between ecology and metabolism is bidirectional and the effect size can be large in both directions. One aspect of this relationship that is missing from the current study is spatial: what are the substrate availability and competitive landscape in the site of colonization?

Manipulating the bile acid pool *in situ* is extraordinarily challenging, especially in the context of a complex community. Our attempts to manipulate the bile acid pool rationally led to unexpected phenotypes, likely due to feedback mechanisms. For example, when we tried to increase production of UDCA in *Cyp2c70/Cyp2a12* double-mutant mice, we instead made the bile acid pool larger, likely because the UDCA served as an FXR antagonist.

Nonetheless, there is reason for optimism. For the first time, by removing and adding strains and genes to a defined community, we are exerting a large effect on the bile acid pool. Although we cannot yet do this rationally, we now have the tools to do it empirically. Hopefully—in the process—we will learn the rules that will make future engineering efforts predictable. Moreover, the combination of a defined community and AAV-CRISPR constructs to mutate host bile acid enzymes is powerful; it affords an unprecedented ability to toggle on and off different components of the bile acid pool, which will help define the role of individual components of the pool in physiology and disease. Future efforts could plausibly lead to therapeutic communities that are relevant to cholestatic disease, IBD, obesity, and enteric infection.

## ACKNOWLEDGMENTS

We are deeply indebted to members of the Fischbach laboratory for helpful discussions and suggestions, especially Gabriel Filsinger and Brian Caliando for cloning support and suggestions, Min Wang and Alice Cheng for microbiology input, Thi Phuong Thao Pham for culturomics, and Maria Prado, Amanda Espinoza and Joanne Au for keeping the laboratory running. We thank Paul Dawson for helpful discussions about bile acid metabolism and physiology. X.Z. is a Fraternal Order of Eagles Fellow of the Damon Runyon Cancer Research Foundation, DRG-2517-24. This work was supported by the Helmsley Foundation, the DOW (Vannevar Bush Faculty Fellowship), NIH (DK101674), NSF (2125383), and the Stanford Microbiome Therapies Initiative.

## AUTHOR CONTRIBUTIONS

X.Z. and M.A.F. conceived the study, X.Z. performed most experiments, including data analysis, community construction, cloning, and method development. X.M. helped with bioinformatics, A.M.W, A.V.C. performed metagenomic sequencing, K.E.J., R.L., M.B., and T.Q.V. designed and constructed AAV-CRISPR constructs. S.H., E.M.L., assisted with gnotobiotic experiments, I.J.G., B.C.D. helped with metabolomics, M.T. helped with cloning, A.Z., K.R.H., M.I, and J.A. helped with community preparation. The manuscript was drafted by X.Z. and M.A.F. and reviewed and edited by X.Z., M.A.F., K.E.J., and T.Q.V.

## COMPETING INTERESTS

M.A.F. is a co-founder of Revolution Medicines and Kelonia, a co-founder and director of Azalea, a member of the scientific advisory boards of the Chan Zuckerberg Initiative, NGM Biopharmaceuticals, and TCG Labs/Soleil Labs, and a science partner at The Column Group.

## SUPPLEMENTARY MATERIALS

## Materials and Methods

Figures s1-s5

Table s1-s4

## MATERIALS AND METHODS

### Bacterial strains and culture conditions

All bacterial strains used in the synthetic community were sourced from the American Type Culture Collection (ATCC), the Leibniz Institute DSMZ–German Collection of Microorganisms and Cell Cultures GmbH (DSMZ), BEI Resources (BEI), and additional sources as specified. Strains were cultured in one of the following media: modified Columbia medium, modified Peptone Yeast Extract Glucose (PYG) medium, Yeast Casitone Fatty Acids (YCFA) medium, modified Gifu medium, modified Baar medium, bifidobacterium medium, ethanoligenens medium, or chopped meat medium. Where necessary, specific nutrients were added to individual cultures to support optimal growth (see **Table S1**).

### Preparation of synthetic communities

For all community culturing, strains were cultured in anaerobic conditions (10% CO_2_, 5% H_2_, and 85% N_2_). For strain purification and storage, each strain was streaked on its respective agar (see **Table S1**), assessed for purity, and a single colony was propagated in 200-300 mL of the corresponding liquid medium. Cultures were harvested at early stationary phase, centrifuged, and resuspended in 25% glycerol prepared in the same growth medium. Aliquots of 300 μL were stored in individual matrix tubes at –80 °C.

To prepare synthetic communities for *in vivo* experiments, frozen stocks of each strain were thawed and propagated in 40 mL of their respective growth medium in 50 mL Falcon tubes for 72 h. Equal volumes of each culture comprising the intended community (including hCom2a and strain dropout communities) were pooled, centrifuged at 4,700 × g for 30 minutes, washed, and resuspended in an equal volume of 25% glycerol in modified Columbia medium. The final synthetic community mixture was aliquoted into 700 μL portions and stored at –80 °C until use.

### Gnotobiotic mouse experiments

Germ-free Swiss-Webster or C57BL/6N male mice (6–8 weeks of age) were originally obtained from Taconic Biosciences (Hudson, NY). Colonies were maintained in gnotobiotic isolators and provided autoclaved chow and water *ad libitum*. All animal procedures were approved by the Stanford University Institutional Animal Care and Use Committee (IACUC).

For colonization, 1.2 mL glycerol stocks of synthetic communities were thawed and mixed thoroughly at room temperature. Mice were orally gavaged with 200 μL of the mixed culture. To ensure robust colonization by all strains, mice were gavaged once daily for two consecutive days in all experiments.

The mice were maintained on a standard diet (LabDiet 5k67; 0.2% Trp) for 4 weeks (unless otherwise stated) before sacrifice (fed *ad libitum*). Mice were euthanized humanely by CO_2_ asphyxiation and, urine, liver, plasma the luminal contents of the small intestine, cecum, and colon were collected at the same time of day and stored at –80 °C until use.

### Metagenomic sequencing

Bacterial isolates and synthetic communities were processed using a unified sequencing workflow. Cell pellets were obtained by centrifugation under anaerobic conditions. Genomic DNA was isolated using the DNeasy PowerSoil HTP Kit (Qiagen), and DNA concentrations were quantified in 384-well plates using the Quant-iT PicoGreen dsDNA Assay Kit (Thermo Fisher). Library preparation was conducted in 384-well format using a miniaturized protocol adapted from the Nextera XT method (Illumina). DNA input was normalized to 0.18 ng/μL using a Mantis liquid handler (Formulatrix). Samples with lower concentrations were not diluted further. Tagmentation, neutralization, and PCR amplification were performed using the Mosquito HTS (TTP Labtech), with a final reaction volume of 4 μL per sample. Custom 12-bp dual unique indexing primers were used during PCR to prevent barcode misassignment associated with patterned flow cells. Libraries were pooled at specified molar ratios and purified with Ampure XP beads (Beckman Coulter) to remove contaminants and size-select fragments between 300 bp and 1.5 kb. Final library pools were assessed for quality and concentration using a Fragment Analyzer (Agilent) and quantitative PCR (Bio-Rad). Sequencing was carried out on either the NovaSeq S4 or NextSeq High Output platforms (Illumina), with 2×150 bp read lengths. We aimed to obtain 5–10 million paired-end reads per bacterial isolate and 20–30 million reads per synthetic community sample.

### Metagenomic mapping

Paired-end sequencing reads were aligned to the hCom2a reference genome database using Bowtie2 (v2.3.7) with the following parameters: --very-sensitive, --maxins 3000, -k 300, --no-mixed, --no-discordant, --end-to-end, and --no-unal. These settings enforced highly sensitive global alignments while suppressing unpaired, discordant, and unaligned reads. The alignment output in SAM format was streamed directly into Samtools (v1.9) for conversion to BAM format, followed by coordinate-based sorting and indexing.

Taking advantage of the fact that every organism in the hCom2 database has been carefully assembled, we used a highly sensitive pipeline, NinjaMap, to translate read alignment in the above BAM file into community composition, including detection of very low abundance organisms (<10^-6^). The complete metagenomic read analysis and the specific read mapping pipeline are available at https://github.com/FischbachLab/nf-ninjamap. To maximize accuracy and mitigate errors arising from imperfect database genomes in this study, we masked regions in NinjaMap based on mis mapped regions identified from the dropout strains in the dropout experiments. Reads that map to these masked regions in the subsequent analyses are assigned directly to the escrow bin.

To ensure high-confidence alignments, only reads with ≥99% identity across their full length were retained. On average, 4.95% of reads (range: 4.10%–8.35%) did not align to the reference. To investigate the source of these unaligned reads, a representative subset was queried against the NCBI nucleotide (nt) database using BLAST+ (v2.11.0) via the ncbi/blast:latest Docker image. The following parameters were applied: -outfmt ’6 std qlen slen qcovs sscinames staxids’, -dbsize 1000000, and -num_alignments 100.

Top BLAST hits were defined as alignments with e-values ≤ 1e–30, percent identity ≥ 99%, and bit scores within 10% of the best hit for each query. Taxonomic summaries were generated using the ktImportTaxonomy script from the Krona toolkit, with default settings. Reads were aggregated by NCBI Taxonomy ID and genus. Most hits corresponded to taxa closely related to members of the synthetic community, while a portion mapped to the mouse genome. These results suggest a negligible level of environmental or cross-sample contamination in our sequencing data.

### Sample preparation for metabolomics

Mouse cecal contents and urine samples were processed for untargeted metabolomics analysis. For cecal samples, wet tissue (20–40 mg) was weighed into 2 mL screw-cap tubes preloaded with six 2.3 mm ceramic beads (RPI research products 9839). An extraction solvent consisting of 40:40:20 methanol: acetonitrile: water (v/v/v), containing 5 μM 4-chlorophenylalanine as an internal standard, was added at a volume of 20 μL per mg of tissue. Samples were homogenized using a QIAGEN TissueLyser II at 25 Hz for 10 minutes. Following homogenization, samples were centrifuged at 18,200 × g for 15 minutes at 4 °C. The resulting supernatants were collected, and 5 μL was injected for liquid chromatography–mass spectrometry (LC-MS) analysis. For mouse urine samples, 5 μL of urine was extracted with 100 μL of methanol containing 5 μM 4-chlorophenylalanine as an internal standard. Samples were incubated on ice for 20 minutes to facilitate protein precipitation, then centrifuged at 18,200 × g for 15 minutes at 4 °C. The supernatant was collected, and 5 μL was injected for LC-MS analysis.

Mouse gallbladder and small intestine samples were extracted for bile acid analysis by LC–MS. Gallbladder contents (2 µL) were extracted in 100 µL methanol containing 5 µM d^4^-cholic acid as an internal standard. Samples were vortexed briefly and centrifuged at 18,200 × g for 10 min at 4 °C. The supernatant was diluted 10-fold in methanol prior to transfer to LC–MS vials for analysis.

For small intestine measurements, in some experiments, only a portion of the small intestine was extracted for bile acid analysis to preserve remaining tissue for parallel metagenomic sequencing. In these cases, regional bile acid measurements were used to assess local bile acid concentrations (reported in µM) rather than total intestinal bile acid pool size (reported in µmol). To measure total bile acid pool, the entire small intestine (including duodenum, jejunum, and ileum) was transferred to a 5 mL tube (Qiagen 990552) and homogenized in 3 mL 40:40:20 methanol: acetonitrile: water (v/v/v) with 5 µM d^4^-cholic acid with 6.5 mm ceramic beads (Omni 19-682). Homogenates were centrifuged at 18,200 × g for 15 min at 4 °C, and the resulting supernatants were diluted 100-fold in methanol before LC–MS analysis. For regional bile acid measurements, approximately 100 mg of small intestine tissue (jejunum) was weighed into a 2 mL microcentrifuge tube and homogenized in 40:40:20 methanol:acetonitrile:water (v/v/v) with internal standards at a 100-fold dilution. After clarification by centrifugation, extracts were transferred directly to LC–MS vials for analysis.

### Liquid chromatography/mass spectrometry

Bile acids were analyzed using an Agilent 6530 quadrupole time-of-flight (Q-TOF) mass spectrometer coupled to an Agilent 1290 Infinity II UPLC system. A dual Agilent Jet Stream electrospray ionization (AJS-ESI) source was used under extended dynamic range (m/z up to 1700). AJS-ESI source settings were as follows: gas temperature, 300 °C; drying gas flow, 7.0 L/min; nebulizer pressure, 40 psig; sheath gas temperature, 350 °C; sheath gas flow, 10.0 L/min; capillary voltage (VCap), 3500 V; nozzle voltage, 1400 V; and fragmentor voltage, 200 V.

Bile acids were separated by a Kinetex C18 column (1.7 μm, 2.1 × 100 mm; Phenomenex). Mobile phase A consisted of 0.05% formic acid in water, and mobile phase B consisted of 0.05% formic acid in acetone. 5 μL of each sample was injected via autosampler. Chromatographic separation was achieved using the following gradient at a flow rate of 0.35 mL/min: 25% B at 0–1 min; linear increase to 75% B at 25 min; ramp to 100% B at 26 min; held at 100% B until 30 min; and re-equilibrated to 25% B at 32 min.

Aromatic amino acids derivatives were analyzed using a Thermo Q Exactive HF Hybrid Quadrupole-Orbitrap mass spectrometer coupled to a Thermo Vanquish UPLC system. MS1 scans were acquired over an m/z range of 60–900 at a resolution of 60,000. Data was collected in centroid mode with a loop count of 4 and an isolation window of 1.2 Da. Chromatographic separation was performed using an Acquity UPLC BEH C18 column (2.1 mm × 100 mm, 1.7 μm particle size, 130 Å pore size; Waters, Milford, MA). A 5 μL aliquot of each sample was injected via autosampler. The mobile phases consisted of water (A) and methanol (B), both containing 0.1% formic acid. Separation was achieved at a flow rate of 0.35 mL/min using the following gradient: 0 min, 0.5% B; 4 min, 70% B; 4.5–5.4 min, 98% B; 5.6 min, re-equilibrated to 0.5% B.

### qPCR

We performed RT-qPCR on RNA extracted from liver samples. RNA was extracted using Direct-zol RNA Miniprep kits (Zymo Research). RNA was quantified using a NanoDrop spectrophotometer (Thermo Fisher). cDNA libraries were generated using the iScript™ cDNA Synthesis Kit (Bio-Rad) or qScript cDNA SuperMix (Quantabio). qPCR was then performed using the qRT-PCR Brilliant III SYBR Master Mix. Relative enrichment was calculated individually for each gene using the ΔΔCt method. Primers used in the study were adopted from previous published studies^30,39^ and list in **Table S4a**. All genes were normalized to either *Tbp* or *36B4*.

### AAV-CRISPR design, cloning and production

Detailed methods for AAV-CRISPR design, cloning, and production have been described previously^15,41,42^, and the AAV-CRISPR constructs utilized here targeting *Cyp2a12* and *Cyp2c70* were recently described and validated^15^. In brief, AAV was used to deliver a complete CRISPR system based on *Staphylococcus aureus* CRISPR/Cas9. Small gRNAs targeting *Cyp2a12* and *Cyp2c70* were cloned into and expressed from the 1313 pAAV-U6-BbsI-MluI-gRNA-SA-HLP-SACas9-HA-OLLAS-spA backbone (Addgene, 109304), a gift from Dr. William Lagor (Baylor College of Medicine)^43^. In all experiments, the control AAV-CRISPR was used, the control AAV-CRISPR constructs were designed by targeting pAAV-U6-BbsI-MluI-gRNA-SA-HLP-SACas9-HA-OLLAS-spA backbone.

For the production of AAV2/8, the triple transfection method followed by cesium chloride gradient was used as previously described^15,44^. Following qPCR titer of the purified virus, aliquots of the AAV were stored until use. For experiments, mice were dosed with 5×10^11^ genome copies per mouse in sterile PBS via intraperitoneal injection. Successful injection was confirmed by observing expected change in bile acid pool following metabolomics analysis.

### *In vitro* bile acid assay

To assess bile acid–metabolizing activity, each strain was first cultured overnight in its respective growth medium. The following day, cultures were diluted to pre-log phase (OD_600_ ≈ 0.2) in 4 mL of the same medium supplemented with 100 μM of the indicated bile acid substrate. Cultures were incubated under anaerobic conditions at 37 °C for the specified time. Following incubation, bile acid extraction was performed. Cultures were acidified to pH 1.0 using 6 N HCl, followed by the addition of 400 μL of ethyl acetate. Samples were vortexed thoroughly, centrifuged, and 200 μL the upper organic layer (ethyl acetate) was carefully collected. The ethyl acetate extract was dried and resuspended in 50% methanol (MeOH) in water. Samples were centrifuged and supernatants were transferred to LC-MS vials for analysis.

### Preparation of *Lactobacillus* competent cells and electroporation

Electrocompetent *Lactobacillus* cells were prepared and transformed as previously described^45–50^, with strain-specific modifications. Overnight cultures were sub-cultured into fresh medium and incubated until reaching an OD_600_ of approximately 0.5 (2–5 hours at 37 °C; strains harboring plasmids were incubated at 30 °C). The subculturing media varied by strain as follows:

- **MRS:** *L. gasseri* ATCC 33323
- **MRS with 1% glycine:** *L. plantarum* WCFS1, *L. plantarum* MAF1421, *L. curvatus* MAF2267
- **MRS with 2% glycine and 10 μg/mL ampicillin:** *L. sakei* MAF2176, *L. salivarius* MAF2147, *L. rhamnosus* ATCC 53103, *L. paracasei* MAF2290

Cells were harvested by centrifugation and washed with ice-cold buffer specific to each strain:

- **10% glycerol:** *L. plantarum* WCFS1, *L. plantarum* MAF1421, *L. curvatus* MAF2267
- **KEB buffer** (0.5 M sucrose, 1 mM MgCl_2_, 7 mM potassium phosphate, pH 7.2): *L. rhamnosus* ATCC 53103
- **HEB buffer** (0.272 M sucrose, 1 mM MgCl_2_, 7 mM HEPES, pH 7.3): *L. sakei* MAF2176, *L. salivarius* MAF2147, *L. paracasei* MAF2290
- **3.5× SMEB buffer** (0.952 M sucrose, 3.5 mM MgCl_2_, pH 7.2): *L. gasseri* ATCC 33323

Cells were resuspended in the corresponding chilled buffer and kept on ice until transformation. Approximately 1–5 μg of plasmid DNA was gently mixed with 50–100 μL of competent cells and transferred into a pre-chilled 1mm or 2 mm electroporation cuvette (Bio-Rad). Electroporation was performed under the following conditions, yielding a typical time constant of approximately 5 ms:

- **1.8 kV in 1mm cuvette:** *L. plantarum* WCFS1, *L. plantarum* MAF1421, *L. curvatus* MAF2267, *L. sakei* MAF2176, *L. paracasei* MAF2290
- **1.8 kV in 2mm cuvette:** *L. gasseri* ATCC 33323, *L. salivarius* MAF2147
- **1.7 kV in 2mm cuvette:** *L. rhamnosus* ATCC 53103

Immediately after electroporation, 1 mL of pre-warmed recovery buffer (0.01 M CaCl₂ and 0.5 M sucrose in MRS) was added to each cuvette. Cells were incubated at 30 °C for 2–3 hours before plating onto MRS agar supplemented with the appropriate antibiotic for selection.

### Disruption of *LP_3536* in *L. plantarum* WCFS1

The gene *LP_3536*, previously identified as encoding a major bile salt hydrolase in *L. plantarum WCFS1*^51^, was targeted for deletion. For all the PCR amplification steps, we used either PrimeSTAR MAX DNA polymerase (Takara Bio) or Q5 High-Fidelity DNA Polymerase (New England Biolabs) according to the manufacturer’s instructions. Primer sequences are shown in (Supplementary Table X). We used a previously reported double-crossover recombination method to construct *L. plantarum* Δ*LP_3536*. In brief, -1kb DNA fragments corresponding to the upstream and downstream regions of the target gene were PCR amplified. The fragment was then re-amplified using primers containing the desired overhangs. The resulting PCR product was assembled into a disruption plasmid pGID023 backbone using Gibson assembly (New England Biolabs) (**Table. S4b**). The resulting plasmid was transformed into *E. coli* DC10B (DH10 Δ*dcm*) by chemical transformation. An *E. coli* clone containing the plasmid was cultivated and plasmid DNA was isolated, purified and verified by DNA sequencing.

The disruption plasmid was introduced into *L. plantarum* WCFS1 via electroporation and plated on MRS agar supplemented with 10 μg/mL erythromycin. Transformants were sub-cultured twice in antibiotic-free MRS medium at 30 °C and then plated on MRS agar with 10 μg/mL erythromycin and incubated at 37 °C to isolate single-crossover integrants. Integration at the target locus was confirmed by genomic PCR. Single-crossover colonies were further passaged in MRS medium at 37 °C without antibiotics for three consecutive days to promote second crossover events. Cultures were diluted and plated on antibiotic-free MRS agar, and resulting colonies were screened for erythromycin sensitivity. Sensitive colonies were subsequently verified by PCR to confirm the successful deletion of *LP_3536*.

### Phage-assisted dsDNA recombineering in *Lactobacillus*

To achieve stable chromosomal integration of exogenous genes in *Lactobacillus* species, we employed a phage-assisted double-stranded DNA (dsDNA) recombineering strategy^47,49^. A helper plasmid encoding the *recT*/*recE* recombinase system under an inducible promoter was first introduced into the *Lactobacillus* strain via electroporation. The Lactobacillus carrying a plasmid was induced to overexpress recT/recE, and were transformed a second time with a linear dsDNA fragment encoding the gene(s) of interest (**Table. S4b**), flanked by ∼1.5 kb homology arms targeting the desired chromosomal integration site. Three helper plasmids were used for different strains. The strains using a pLH plasmid (pLH01 and pLHZ22a) were induced by 100 ng/ ml peptide MAGNSSNFIHKIKQIFTHR for 2 hrs before second electroporation. Strains using a pAD3a plasmid were induced by 500 ng/ml anhydrotetracycline instead.

Strain-specific recombineering helper plasmid were used as follows:

- **pLH01:** L. plantarum WCFS1, *L. plantarum* MAF1421
- **pLHZ22a:** *L. salivarius* MAF2147, *L. rhamnosus* ATCC 53103
- **pAD3a:** *L. curvatus* MAF2267, *L. sakei* MAF2176, *L. paracasei* MAF2290, *L. gasseri* ATCC 33323

pLH01 is a previously described helper plasmid optimized for use in *Lactobacillus plantarum* species^48^. pLHZ22a was derived from pLH01 by replacing the pWV01 origin with a dual-origin system consisting of *E. coli* p15A and gram-positive pAMβ1, enabling replication in both *E. coli* and diverse *Lactobacillus* species. Additionally, the *L. plantarum*–derived *recT/recE* in pLH01 was replaced with a more efficient *recT/recE* pair from *Clostridium strain FS.41*, recently reported to enhance recombination efficiency in closely related lactic acid bacteria^49^. Both pLH01 and pLHZ22a were cloned and propagated in *E. coli DC10B*. Pad3a is a helper plasmid that utilizes an alternative anhydrotetracycline (aTc)-inducible system and was cloned and amplified in *Lactococcus lactis* NZ9000. This plasmid was used for recombineering in strains where *L. lactis* served as a more compatible cloning host.

To generate dsDNA templates for recombineering, different bile acid–modifying enzyme sequences flanked by homology arms were prepared as follows. For *bsh*, the *L. plantarum* gene *LP_3536* was placed under the control of a previously reported strong constitutive promoter^34^. For the urso bile acid pathway, codon-optimized sequences of 7α-hydroxysteroid dehydrogenase (7α-HSDH, Uniprot Q03906) and 7β-hydroxysteroid dehydrogenase (7β-HSDH, Uniprot A7B4V1) were assembled under the same promoter. A ribosome binding site upstream of the 7β-HSDH gene was designed using an RBS calculator (Salis Lab). The complete expression cassette—including promoter, *bsh*, and/or 7α-HSDH, RBS, and 7β-HSDH—was flanked by ∼1.5 kb homology arms corresponding to the target chromosomal integration site. The full construct was cloned into a pSC101-based plasmid and sequence-verified prior to use as a template for recombineering.

