## Supplementary information for "Ecology and engineering to modify the bile acid output of a defined microbial community"

**This PDF file includes:**

Supplementary text
Figures s1-s5
Table s1-s4

### SUPPLEMENTARY TEXT

#### *Ruminococcus gnavus* drives urso-bile acid production in Δ*CspA*

In mice colonized with a community deficient in *Clostridium sp. ATCC 29733* (Δ*CspA*), we observed a selective increase in urso (7β) bile acids. The cecal levels of ursodeoxycholic acid (UDCA), ursocholic acid (7β-CA), 7-oxoCA and the UDCA metabolite β-muricholic acid were each increased by 2-fold (**fig. s1a, s1e-f**). Since epimerization of the 7α-hydroxyl group to form urso bile acids competes with 7α-dehydroxylation to form DCA and LCA (**Fig. 1a**), these findings suggest a metabolic shift toward the former and away from the latter in Δ*CspA*. Notably, this effect was restricted to the cecum, as TUDCA levels in the small intestine remained unchanged (**fig. s1g**).

To identify the microbial basis of the urso–bile acid increase in Δ*CspA*, we performed metagenomic sequencing of cecal contents and analyzed strain-level abundances using NinjaMap. Overall community composition was largely unchanged between the parent and dropout communities, with only a small subset of strains showing substantial abundance shifts. In the Δ*CspA* community, we observed a three-fold increase in *Ruminococcus gnavus*, a well-known producer of urso-bile acids[1] (**fig. s1h**). To assess whether other members of hCom2 might contribute to this activity, we performed a BLAST search for 7β-hydroxysteroid dehydrogenase (7β-HSDH) homologs. Aside from *R. gnavus*, only *Collinsella aerofaciens* was found to harbor a 7β-HSDH gene (**fig. s1i**). However, *C. aerofaciens* was below the detection limit in both control and Δ*CspA* communities (**fig. s1h**). Based on these findings, we conclude that the increased abundance of *R. gnavus* is responsible for the selective production of urso bile acids observed in the Δ*CspA* dropout.

#### *Bacteroides ovatus* drives increased secondary bile acids production in Δ*Ro*

Among the five dropout communities with increased LCA abundance, four showed a concomitant increase of *L. plantarum* abundance, consistent with its likely contribution to the phenotype. In contrast, in the fifth community (Δ*Ro*), *L. plantarum* abundance decreased. In this case, *B. ovatus* was the only strain exhibiting a significant increase in abundance (**fig. s2c-d**).

Notably, stochastic variation in *B. ovatus* colonization within the hCom2a control group provided an internal validation of its role. Four mice exhibiting lower *B. ovatus* abundance (**fig. s2e**) exhibited a significant reduction in taurine-conjugated secondary bile acids (TDCA, TLCA) and the total size of the small intestinal bile acid pool (**fig. s2f-k**). These data suggest that *B. ovatus*—rather than *L. plantarum*—is responsible for the observed expansion of the secondary bile acid pool in ΔRo.

Notably, *B. ovatus,* like *L. plantarum*, encodes bile salt hydrolase[2], indicating that distinct ecological perturbations may promote different *bsh*-harboring strains to bloom, converging on a similar functional outcome.

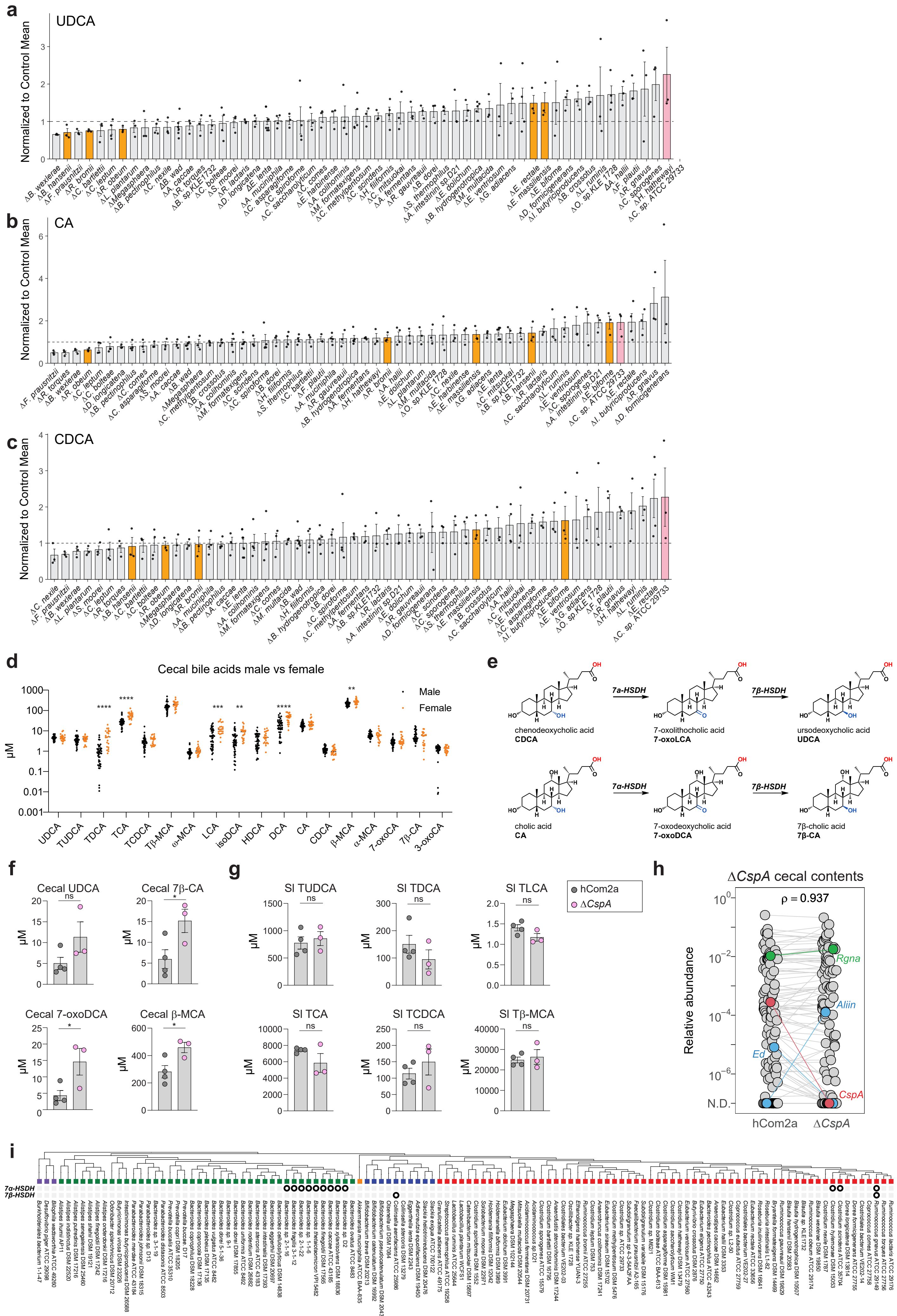
a
UDCA
Normalized to Control Mean
b
CA
Normalized to Control Mean
c
CDCA
Normalized to Control Mean
d
Cecal bile acids male vs female
μM
Male
Female
UDCA
TUDCA
TDCA
TCA
TCDCA
Tβ-MCA
α-MCA
LCA
isoDCA
HDCA
DCA
CA
CDCA
β-MCA
α-MCA
7-oxoCA
7β-CA
3-oxoCA
e
chenodeoxycholic acid
CDCA
7α-HSDH
7-oxolithocholic acid
7-oxoLCA
7β-HSDH
ursodeoxycholic acid
UDCA
cholic acid
CA
7α-HSDH
7-oxodeoxycholic acid
7-oxoDCA
7β-HSDH
7β-cholic acid
7β-CA
f
Cecal UDCA
Cecal 7β-CA
Cecal 7-oxoDCA
Cecal β-MCA
μM
g
SI TUDCA
SI TDCA
SI TLCA
SI TCA
SI TCDCA
SI Tβ-MCA
hCom2a
ΔCspA
h
ΔCspA cecal contents
ρ = 0.937
Relative abundance
N.D.
Rgna
Aliin
Ed
CspA
hCom2a
ΔCspA
i
7α-HSDH
7β-HSDH

**Figure s1: A single-strain dropout screen reveals communities that confer a larger bile acid pool on mice. Related to Figure 1 and supplementary text.** (**a-c**) Waterfall plots depicting the levels of (**a**) ursodeoxycholic acid (UDCA), (**b**) cholic acid (CA), and (**c**) chenodeoxycholic acid (CDCA) in each dropout communities, normalized to the corresponding control group. (n=3-6 per condition). (**d**) Consistency of cecal bile acid profiles between male and female. Cecal bile acid concentrations in male and female hCom2a colonized Swiss-Webster mice. (n=48 for male, n=16 for female) (**e**) Schematic of the enzymatic conversion of primary bile acids (CA and CDCA) into their 7β-epimers. This pathway starts with the oxidation of the 7a-hydroxyl group by 7α-HSDH to generate the intermediate 7-oxo species (7-oxoDCA, and 7-oxoLCA). Next, 7-oxoDCA and 7-oxoLCA were subsequently reduced by 7β-HSDH to form 7β-CA and UDCA. (**f-g**) Selective accumulation of urso-bile acids in Δ*CspA* in cecal contents, but not small intestine. 7β-bile acid species and intermediates levels in the (**e**) cecum and (**f**) small intestine were measured by targeted LC-MS. (n=4 for hCom2a and n=3 for Δ*CspA*). (**h**) The removal of *CspA* triggers a significant expansion of *R. gnavus*. Each dot is an individual strain; the collection of dots in a column represents the community with median values selected from 3-4 mice co-housed in one cage. The strain dropped out is highlighted in red; strains whose relative abundances change significantly are highlighted in blue (student t test $p<0.05$, fold change >10), and *Ruminococcus gnavus* (*Rgna*) is highlighted in green. (**i**) *In silico* analysis of hCom2a strains encoding 7α-HSDH, 7β-HSDH, which mediate bile acid 7α-epimerization. For all panels, data are shown as mean +/- SEM. Statistical significance in panel (**d**) was assessed using a two-tailed student's t-test with False Discovery Rate (FDR) correction: $q < 0.05$ (*), $q < 0.01$ (**), $q < 0.001$ (***), $q < 0.0001$ (****). In other figure panels, statistical significance was determined using two-tailed Student's t-test; $p < 0.05$ (*), $p < 0.01$ (**), $p < 0.001$ (***).

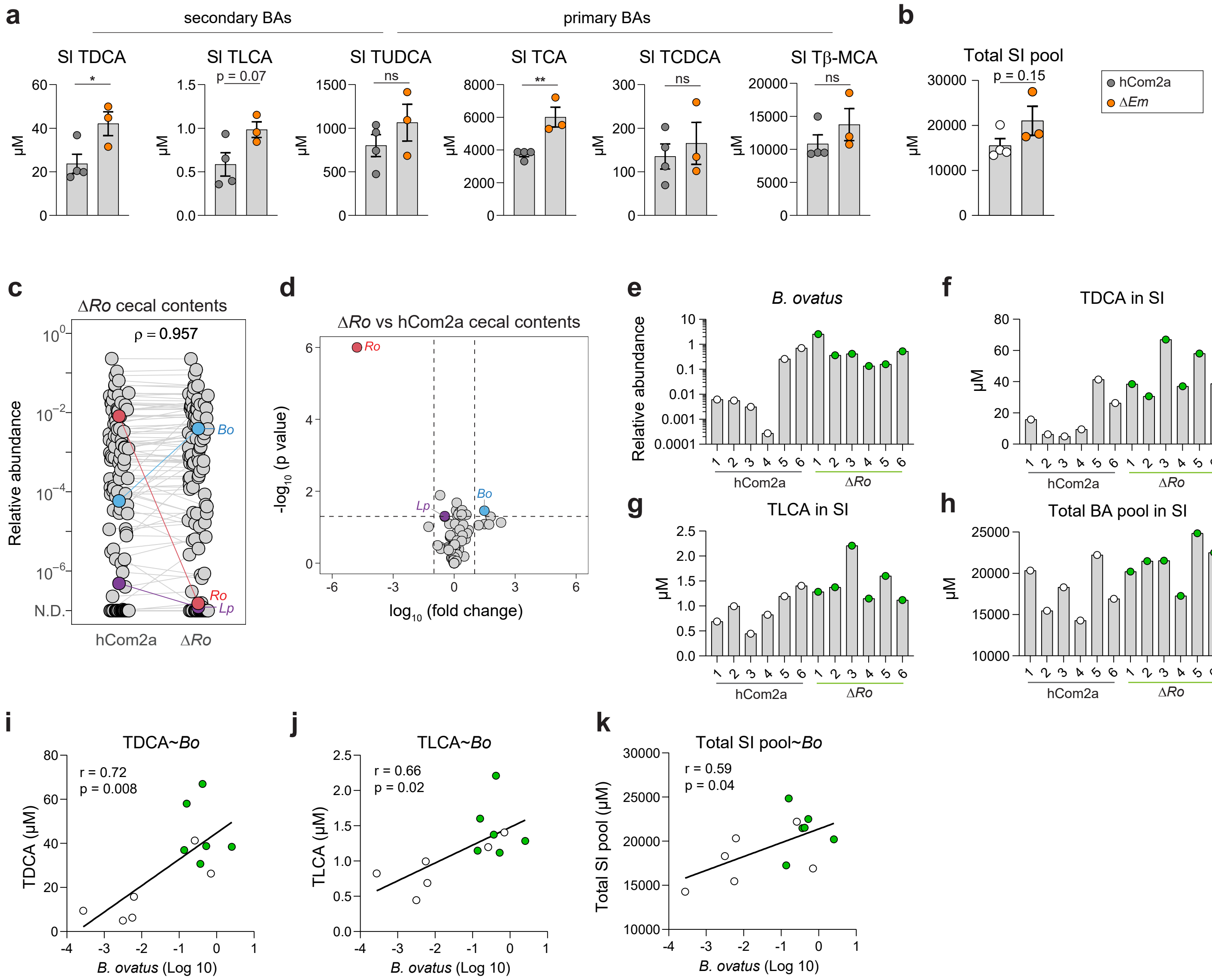

a
secondary BAs
primary BAs
SI TDCA
SI TLCA
SI TUDCA
SI TCA
SI TCDCA
SI Tβ-MCA
p = 0.07
ns
**
ns
ns
*
μM
b
Total SI pool
p = 0.15
hCom2a
ΔEm
c
ΔRo cecal contents
ρ = 0.957
Relative abundance
N.D.
Bo
Ro
Lp
hCom2a
ΔRo
d
ΔRo vs hCom2a cecal contents
-log10 (p value)
log10 (fold change)
e
B. ovatus
Relative abundance
f
TDCA in SI
g
TLCA in SI
h
Total BA pool in SI
i
TDCA~Bo
r = 0.72
p = 0.008
TDCA (μM)
B. ovatus (Log 10)
j
TLCA~Bo
r = 0.66
p = 0.02
TLCA (μM)
k
Total SI pool~Bo
r = 0.59
p = 0.04
Total SI pool (μM)

**Figure s2: Identifying the strains driving an enlarged bile acid pool in ∆*Ro*. Related to Figure 2 and supplementary text.** (**a**) ∆*Em*-colonized mice have an increased bile acid pool size in the small intestine. Small intestinal contents were measured by targeted LC-MS. (n=4 for hCom2a and n=3 for ∆*Em*). (**b**) Total small intestinal bile acid pool size (representing the sum of all measured bile acid species) in mice colonized with hCom2a and ∆*Em* (n=4 for hCom2a and n=3 for ∆*Em*). (**c**) Median relative abundances of hCom2a and ∆*Ro* in cecal contents. Each dot is an individual strain; the collection of dots in a column represents the community with median values selected from 3-4 mice co-housed in one cage. The strain being dropped out is highlighted in red; the strain whose relative abundances change significantly (*Bacteroides ovatus, Bo*) is highlighted in blue (student t test $p<0.05$, fold change >10), and *Lactobacillus plantarum* (*Lp*) is shown in purple. (**d**) Volcano plot comparing the magnitude and significance of strain-level shifts in the cecum between hCom2a and ∆*Ro* mice. (**e**) Relative abundance of *B. ovatus* in individual mice. In hCom2a group, mouse 1-4 exhibited lower *B. ovatus* relative abundance compared to mouse 5-6. (**f-h**) Concentrations of (**f**) TDCA, (**g**) TLCA, (**h**) total bile acid pool in small intestinal contents of each individual mice colonized with hCom2a and ∆*Ro*. (**i-k**) Correlations between TDCA (**h**), TLCA (**i**), or total bile acid pool (**k**) and *B. ovatus* relative abundances in mice colonized with hCom2a and ∆*Ro,* each dot is an individual mouse. White dots represent hCom2a-colonized mice while green dots represent ∆*Ro* colonized mice. All graphs show mean +/- SEM. Statistical significance in panel (**a-b**) was determined using two-tailed student's t-test, statistical significance in panel (**i-k**) was assessed by linear regression analysis (two-tailed test for non-zero slope). $p < 0.05$ (*), $p < 0.01$ (**).

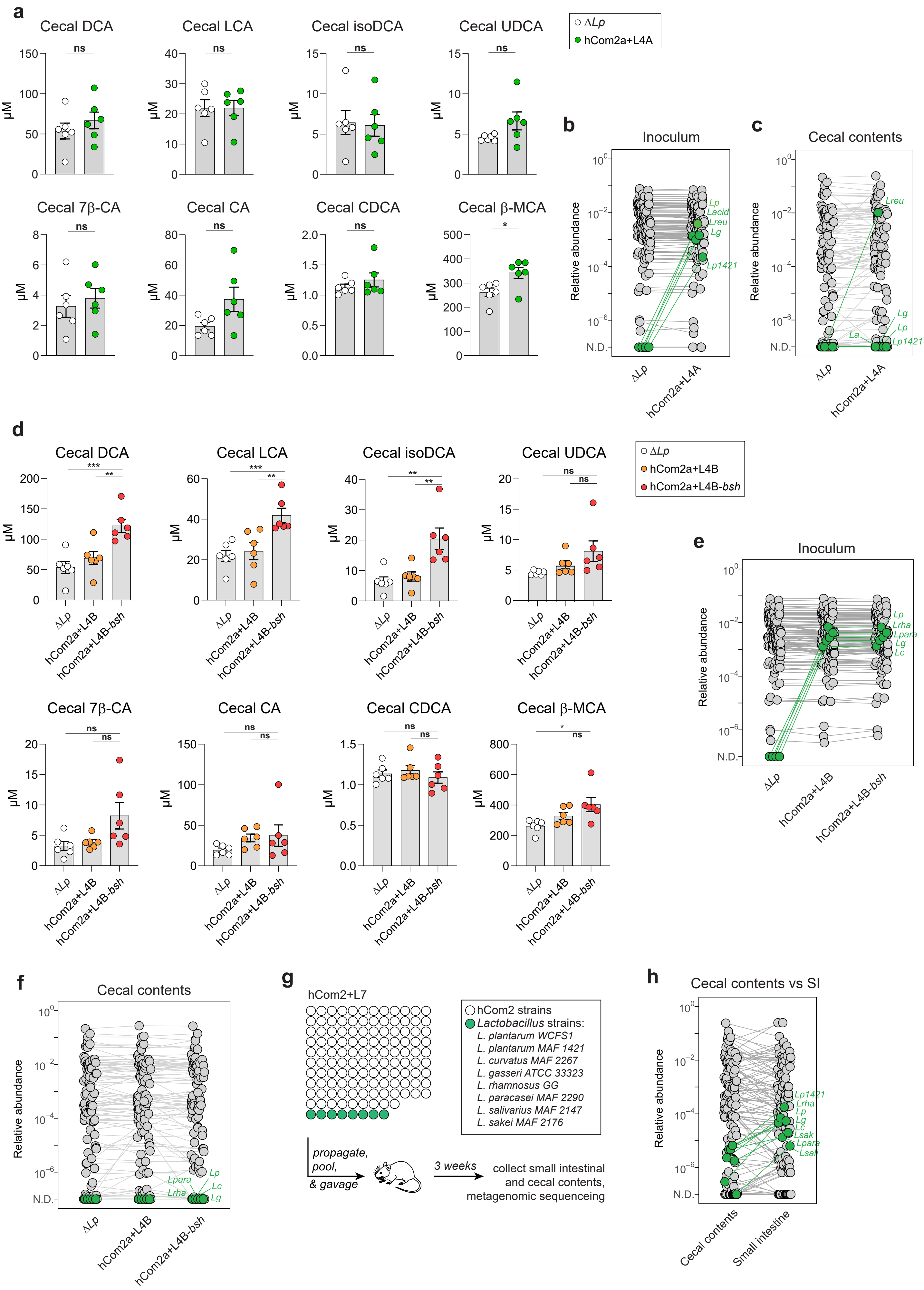
a
Cecal DCA
Cecal LCA
Cecal isoDCA
Cecal UDCA
Cecal 7β-CA
Cecal CA
Cecal CDCA
Cecal β-MCA
μM
ns
*
ΔLp
hCom2a+L4A
b
Inoculum
c
Cecal contents
Relative abundance
N.D.
Lp
Lacid
Lreu
Lg
Lp1421
La
d
e
hCom2a+L4B
hCom2a+L4B-bsh
Lrha
Lpara
Lc
f
g
hCom2+L7
hCom2 strains
Lactobacillus strains:
L. plantarum WCFS1
L. plantarum MAF 1421
L. curvatus MAF 2267
L. gasseri ATCC 33323
L. rhamnosus GG
L. paracasei MAF 2290
L. salivarius MAF 2147
L. sakei MAF 2176
propagate, pool, & gavage
3 weeks
collect small intestinal and cecal contents, metagenomic sequenceing
h
Cecal contents vs SI
Cecal contents
Small intestine
Lsak
Lsal

**Figure s3: Community-level *bsh* engineering increases secondary bile acid production. Related to Figure 3.** (**a**) Targeted metabolomics of cecal contents. The addition of bsh-expressing *Lactobacillus* (hCom2a+L4A) did not increase secondary bile acid levels compared to Δ*Lp* (n=6 per group). (**b**) Median relative abundances of Δ*Lp* and hCom2a+L4A inoculum. Each dot is a strain; *Lactobacillus* strains were highlighted in green. (**c**) Median relative abundances of Δ*Lp* and hCom2a+L4A in cecal contents. (**d**) Targeted metabolomics of cecal contents. Mice colonized with hCom2a+L4B-*bsh* had higher levels of secondary bile acid levels than both control groups (n=6 per group). (**e**) Median relative abundances of Δ*Lp*, hCom2a+L4B, and hCom2a+L4B-*bsh* inoculum. (**f**) Median relative abundances of cecal contents colonized with Δ*Lp,* hCom2a+L4B, and hCom2a+L4B-*bsh*. (**g**) Schematic for experimental design: Germ-free C57BL/6 mice were colonized with Δ*Lp,* or Δ*Lp* with eight additional *Lactobacillus* strains (hCom2a+L7) for 3 weeks. Small intestine and cecal contents were collected for metagenomic sequencing to assess preferential colonization of *Lactobacillus* in the small intestine. (**h**) *Lactobacillus* selectively colonizes the small intestine. Median relative abundances of cecal contents compared to small intestinal contents in hCom2a+L7 colonized mice. All graphs show mean +/- SEM. Statistical significance was determined using two-tailed Student's t-test; $p < 0.05$ (*), $p < 0.01$ (**), $p < 0.001$ (***).

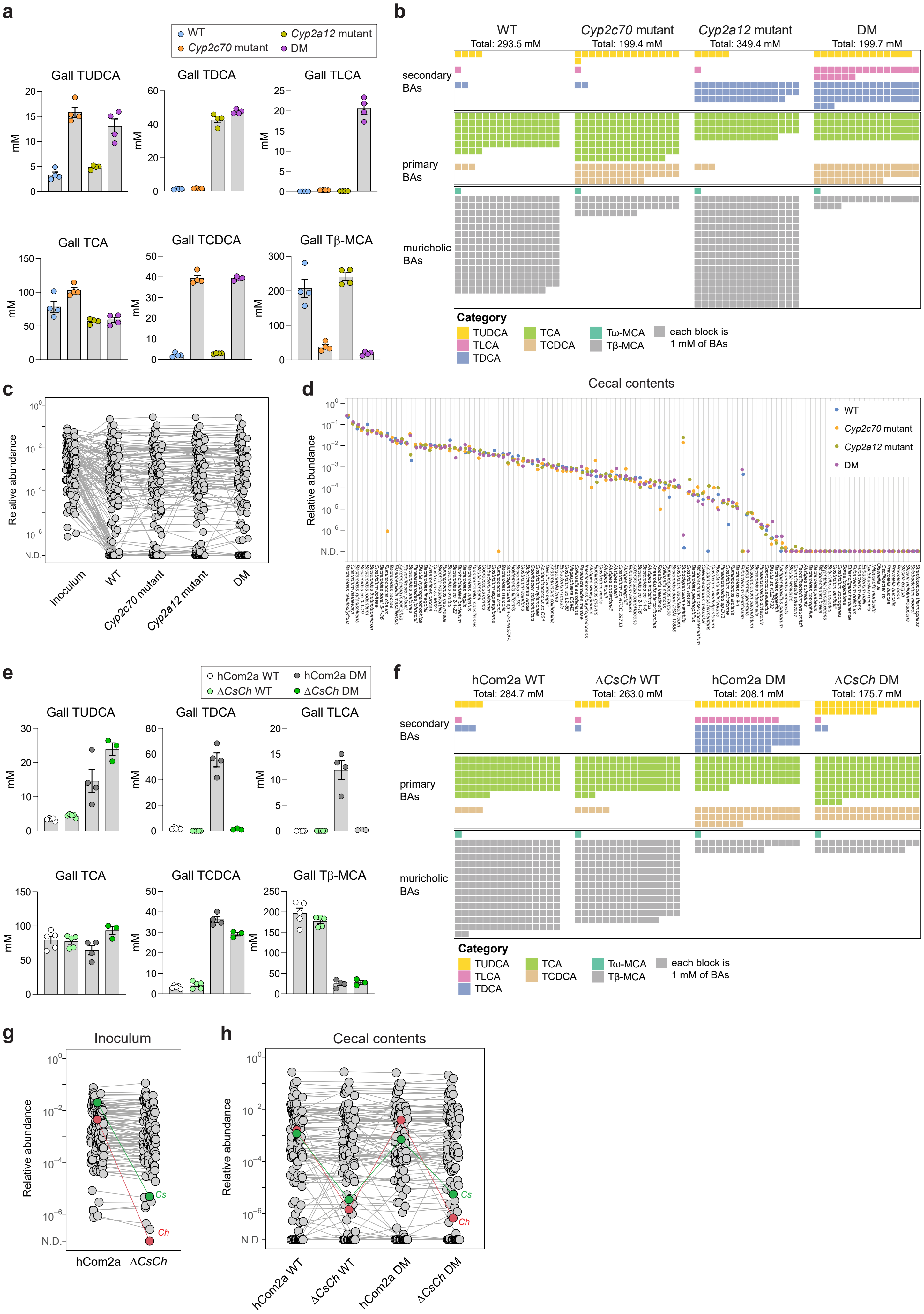

a
WT
Cyp2c70 mutant
Cyp2a12 mutant
DM
Gall TUDCA
Gall TDCA
Gall TLCA
Gall TCA
Gall TCDCA
Gall Tβ-MCA
mM
b
WT
Total: 293.5 mM
Cyp2c70 mutant
Total: 199.4 mM
Cyp2a12 mutant
Total: 349.4 mM
DM
Total: 199.7 mM
secondary BAs
primary BAs
muricholic BAs
Category
TUDCA
TLCA
TDCA
TCA
TCDCA
Tω-MCA
Tβ-MCA
each block is 1 mM of BAs
c
Relative abundance
N.D.
Inoculum
WT
Cyp2c70 mutant
Cyp2a12 mutant
DM
d
Cecal contents
Relative abundance
N.D.
WT
Cyp2c70 mutant
Cyp2a12 mutant
DM
e
hCom2a WT
ΔCsCh WT
hCom2a DM
ΔCsCh DM
Gall TUDCA
Gall TDCA
Gall TLCA
Gall TCA
Gall TCDCA
Gall Tβ-MCA
mM
f
hCom2a WT
Total: 284.7 mM
ΔCsCh WT
Total: 263.0 mM
hCom2a DM
Total: 208.1 mM
ΔCsCh DM
Total: 175.7 mM
secondary BAs
primary BAs
muricholic BAs
Category
TUDCA
TLCA
TDCA
TCA
TCDCA
Tω-MCA
Tβ-MCA
each block is 1 mM of BAs
g
Inoculum
Relative abundance
N.D.
Cs
Ch
hCom2a
ΔCsCh
h
Cecal contents
Relative abundance
N.D.
Cs
Ch
hCom2a WT
ΔCsCh WT
hCom2a DM
ΔCsCh DM

**Figure s4: Characterizing microbial and host effects on the bile acid pool. Related to Figure 4.** (**a**) Targeted metabolomic analysis of gallbladder contents. The bile acid pool in hCom2a-colonized mice is altered profoundly by mutating *Cyp2a12*, *Cyp2c70*, or both, and in the double mutant it more closely resembles the human bile acid pool (n=4 per group). (**b**) A waffle plot showing a visual representation of the bile acid pool in the mice from the experiment schematized in (**a**). Each block corresponds to 1 mM of bile acids. (**c**) Median relative abundances of the inoculum and cecal contents from hCom2a colonized mice across four different genetic background: WT, *Cyp2a12* mutant, *Cyp2c70* mutant, and double mutant (DM). (**d**) Rank-abundance curve of hCom2a strains in the cecal contents. Most strains don't change in relative abundances when host bile acid pool changes, with a few exceptions. (**e**) Targeted metabolomic analysis of gallbladder contents in hCom2a or Δ*CsCh* colonized mice in WT or *Cyp2a12/Cyp2c70* KO mice. Figures show different total taurine-conjugated bile acid levels in the gallbladder in different groups of mice. (**f**) Waffle plot represents the gallbladder bile acid pool in hCom2a or Δ*CsCh* colonized mice in WT or *Cyp2a12/Cyp2c70* KO mice. Each block corresponds to 1 mM of bile acids. (**g**) Median relative abundances of the inoculum hCom2a and Δ*CsCh*. Each dot is an individual strain; the collection of dots in a column represents the community with median values for each strain. *Clostridium scindens* (*Cs*) and *Clostridium hylemonae* (*Ch*) were highlighted in red and green, respectively. (**h**) Median relative abundances of cecal contents from mice colonized with hCom2a and Δ*CsCh*. All graphs show mean +/- SEM. Statistical significance was determined using two-tailed Student's t-test; $p < 0.05$ (*).

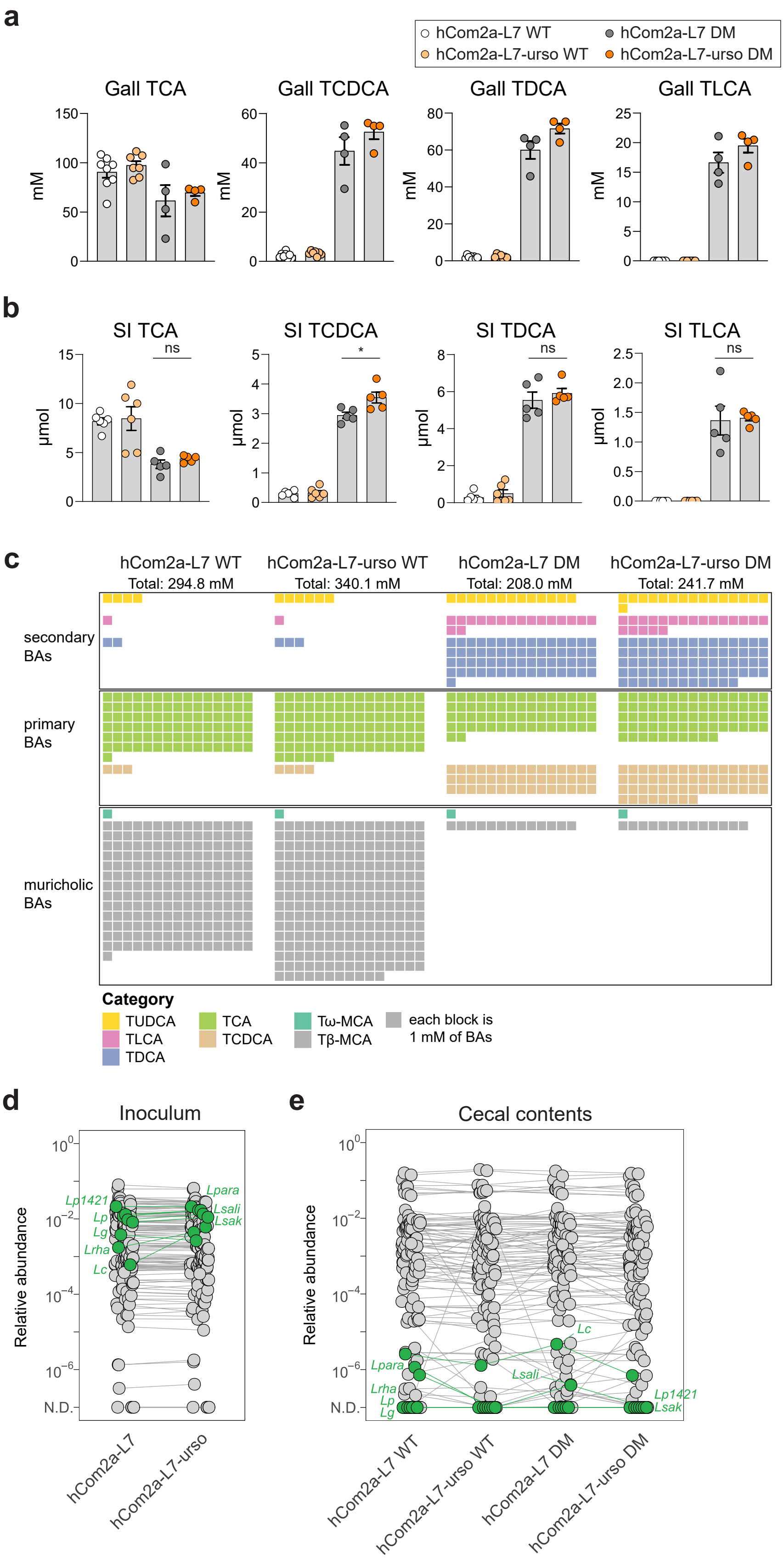
a
hCom2a-L7 WT
hCom2a-L7 DM
hCom2a-L7-urso WT
hCom2a-L7-urso DM
Gall TCA
Gall TCDCA
Gall TDCA
Gall TLCA
mM
b
SI TCA
SI TCDCA
SI TDCA
SI TLCA
µmol
ns
*
c
hCom2a-L7 WT
Total: 294.8 mM
hCom2a-L7-urso WT
Total: 340.1 mM
hCom2a-L7 DM
Total: 208.0 mM
hCom2a-L7-urso DM
Total: 241.7 mM
secondary BAs
primary BAs
muricholic BAs
Category
TUDCA
TLCA
TDCA
TCA
TCDCA
Tω-MCA
Tβ-MCA
each block is 1 mM of BAs
d
Inoculum
Relative abundance
Lp1421
Lp
Lg
Lrha
Lc
Lpara
Lsali
Lsak
N.D.
hCom2a-L7
hCom2a-L7-urso
e
Cecal contents
Relative abundance
Lpara
Lrha
Lp
Lg
Lc
Lsali
Lp1421
Lsak
N.D.
hCom2a-L7 WT
hCom2a-L7-urso WT
hCom2a-L7 DM
hCom2a-L7-urso DM

**Figure s5: Engineering the pool toward urso bile acids. Related to Figure 5.** (**a**) Targeted metabolomics of gallbladder contents. Concentrations of TCA, TCDCA, TDCA, and TLCA in gallbladder were quantified by targeted LC-MS based metabolomics (n=8 for hCom2a WT, n=7 for hCom2a-urso WT, n=4 for DM groups). (**b**) Same as (**a**), for small intestinal contents. (n=6 for WT groups, n=5 for DM groups). (**c**) A waffle plot showing a visual representation of the bile acid pool in the gallbladder of mice from the experiment schematized in (**a**). Each block corresponds to 1 mM of bile acids. (**d**) Median relative abundances of the hCom2a-L7 and hCom2a-L7-urso inoculum. Each dot is an individual strain; the collection of dots in a column represents the community with median values for each strain. Added *Lactobacillus* strains are highlighted in green. (**e**) Median relative abundances of cecal contents from mice colonized with hCom2a-L7 and hCom2a-L7-urso. All graphs show mean +/- SEM. Statistical significance was determined using two-tailed Student's t-test; $p < 0.05$ (*).
